# Dual prognostic role for 2-oxoglutarate oxygenases in ten diverse cancer types: Implications for cell cycle regulation and cell adhesion maintenance

**DOI:** 10.1101/442947

**Authors:** Wai Hoong Chang, Donall Forde, Alvina G. Lai

## Abstract

**Objectives:** Tumor hypoxia is associated with metastasis and resistance to chemotherapy and radiotherapy. Genes involved in oxygen-sensing are clinically relevant and have significant implications on prognosis.

**Methods:** We identified of two signatures, signature 1 (good prognosis) and signature 2 (adverse prognosis), each consisting of 5 genes using three pancreatic cancer cohorts (n=681). We validated the signatures’ performance in predicting survival in ten cancers using Cox regression and receiver operating characteristic (ROC) analyses.

**Results:** Signature 1 and signature 2 were associated with good and poor overall survival respectively. Prognosis of signature 1 in 8 cohorts representing 6 cancers (n=2,627): bladder (hazard ratio [HR]=0.68, P=0.039), papillary renal cell (HR=0.35, P=0.013), liver (HR=0.64, P=0.033 and HR=0.49, P=0.025), lung (HR=0.66, P=0.014) and pancreatic (HR=0.42, P<0.001 and HR=0.64, P=0.04) and endometrial (HR=0.40, P<0.001). Prognosis of signature 2 in 12 cohorts representing 9 cancers (n=4,134): bladder (HR=1.46, P=0.039), cervical (HR=1.97, P=0.035), head and neck (HR=1.39, P=0.038), renal clear cell (HR=1.47, P=0.012), papillary renal cell (HR=3.89, P=0.0015), liver (HR=5.10, P<0.0001 and HR=2.26, P<0.001), lung (HR=1.54, P=0.011), pancreatic (HR=2.09, P=0.002, HR=1.46, P=0.018, and HR=1.99, P<0.0001) and stomach (HR=1.78, P=0.004). Multivariate Cox regression confirmed independent clinical relevance of signatures in these cancers. ROC analyses confirmed superior performance of signatures to current tumor staging benchmarks. *KDM8* is a potential tumor suppressor downregulated in liver and pancreatic cancers and is an independent prognostic factor. *KDM8* expression negatively correlated with cell cycle regulators. Low *KDM8* in tumors was associated with loss of cell adhesion phenotype through *HNF4A* signaling.

**Conclusions:** Pan-cancer signatures of oxygen-sensing genes used for risk assessment in 10 cancers (n=6,761) could guide individualized treatment plans.

## Introduction

Solid tumors face considerable demands in maintaining oxygen availability due to their unique vasculature systems[1]. Rapid neoplastic cell proliferation and overexpression of angiogenic factors leading to the formation of disorganized blood vessels result in insufficient oxygen supply to tumor cells[2,3]. Hence, there is a requirement for tumors to evolve systems that detect changes in oxygen homeostasis. The discovery of hypoxia-inducible factor (HIF) as one of the key oxygen-sensing gene represents a quantum leap forward in tumor biology[4,5]. HIF has co-evolved with the need for more sophisticated oxygen delivery system but intriguingly, it is found in primitive animals that lack oxygen-delivery structures altogether[6]. Its discovery has led to the development of drugs used to treat anemia and cancer[7,8].

In addition to HIF, 2-oxoglutarate (2OG) oxygenases represent another family of oxygen-sensing proteins. As suggested by its name, this group of enzymes has an absolute requirement for molecular oxygen. They catalyze a range of oxidative modifications and their activities are affected by nutrient and oxygen availability[9], both of which are altered within the tumor microenvironment. Several members from this gene family have been implicated in cancer. *Ten-Eleven Translocation 2* (*TET2*) is frequently found to be mutated in leukemia[10] and other solid malignancies[11]. JmjC lysine demethylases (*KDMs*) epigenetically regulate cancer-related genes[12] and inactivating mutations are observed in multiple cancers; glioblastoma/neural, breast, colorectal and renal[13,14].

We hypothesize that 2OG oxygenases expression could help predict prognosis in solid malignancies that are characteristically oxygen-deprived. Additionally, we hypothesize this would be applicable to different types of cancers as they share a uniform need to overcome hypoxia for survival. Starting from an initial set of 61 2OG oxygenases, we developed two prognostic gene signatures, each consisting of a minimal 5 genes, that could facilitate risk stratification and predict overall survival in cancer patients. We confirmed their prognostic performance through a multi-cohort pan-cancer validation process.

## Methods

Detailed methods are available in supplementary information.

## Results

### Multi-cohort analyses revealed two prognostic 2OG oxygenases signatures

We analyzed the prognostic significance of 61 2OG oxygenase genes in 25 cancers (n = 19,781) from The Cancer Genome Atlas[15](Table S1 & S2). Prognostic genes were defined as those whose expression levels were significantly correlated with patients’ overall survival. Since the highest number of prognostic genes (29/61) was observed in pancreatic cancer (n=178) and it is one of the most difficult to treat cancers, two additional pancreatic cancer cohorts from the International Cancer Genome Consortium [ICGC][16] (n=269 and n=234) were used in combination as training cohorts (Fig.1A). We defined two gene signatures; signature 1 and signature 2 as favorable and unfavorable prognostic factors respectively, by taking into consideration genes that were significant in univariate Cox analyses in two out of three pancreatic cancer cohorts (Fig.1A,1B). Signature 1 constitutes 5 genes: *KDM8, KDM6B, P4HTM, ALKBH4* and *ALKBH7*. Likewise, 5 genes make up signature 2: *KDM3A, P4HA1, ASPH, PLOD1* and *PLOD2* (Fig.1A).

**Figure 1.**
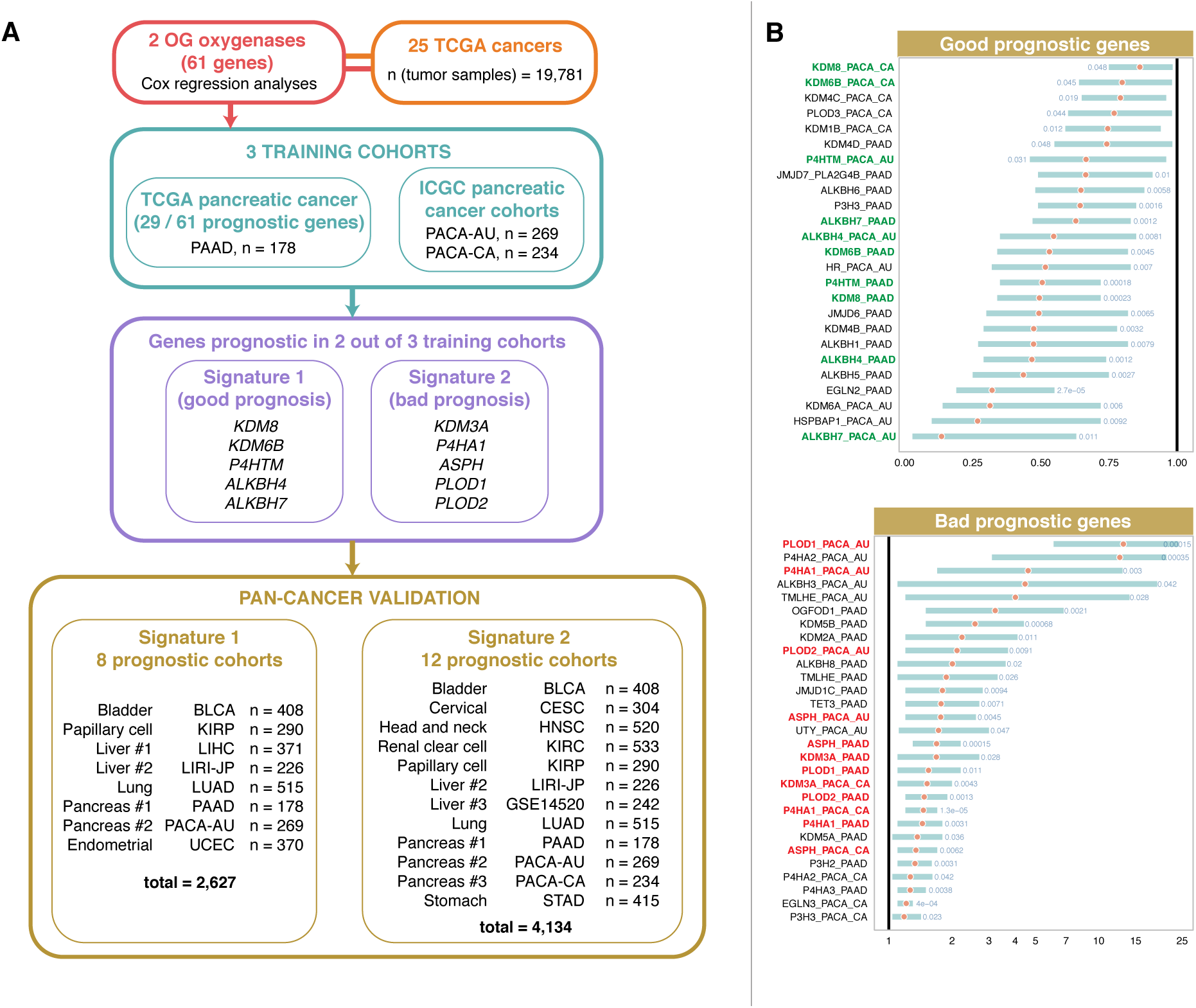
Schematic diagram of the study design and development of signatures derived from oxygen-sensing genes. **(A)** Three pancreatic cancer cohorts were used to define both signatures 1 and 2. Genes found to be prognostic in Cox regression analysis in two out of three pancreatic cancer cohorts were included in signatures 1 and 2. Signature 1 is a marker of good prognosis and consists of 5 genes (*KDM8, KDM6B, P4HTM, ALKBH4* and *ALKBH7*). Signature 2 is a marker of adverse prognosis and consists of 5 genes (*KDM3A, P4HA1, ASPH, PLOD1* and *PLOD2*). Prognosis of both signatures were further confirmed in 10 cancer types using Kaplan-Meier, Cox regression and receiver operating characteristic analyses. **(B)** Forest plots of prognostic genes found to be significant by Cox proportional hazards analysis in pancreatic cancer cohorts abbreviated as PAAD, PACA-AU and PACA-CA. Genes were separated into two groups, good and bad prognostic genes. Hazard ratios (HR) were denoted as red circles and turquoise bars represent 95% confidence interval. Significant Wald test P values were indicated in blue. Y-axes represent gene symbols followed by cohort abbreviations. Signature 1 genes were marked in green. Signature 2 genes were marked in red. Full description of cancers is listed in Table S1.

Patients were median-dichotomized based on mean expression scores of signatures 1 and 2 genes. Patients with high expression of signature 1 genes had significantly better survival in 6 cancers: bladder (hazard ratio [HR]=0.68, P=0.039), papillary renal cell (HR=0.35, P=0.013), liver (HR=0.64, P=0.033 and HR=0.49, P=0.025), lung (HR=0.66, P=0.014), pancreatic (HR=0.42, P<0.001 and HR=0.64, P=0.04) and endometrial (HR=0.40, P<0.001); consistent with the fact that signature 1 is a marker of good prognosis (Fig.2A; Table S3). In contrast, patients with high expression of signature 2 genes had significantly worse prognosis in 9 cancers: bladder (HR=1.46, P=0.039), cervical (HR=1.97, P=0.035), head and neck (HR=1.39, P=0.038), renal clear cell (HR=1.47, P=0.012), papillary renal cell (HR=3.89, P=0.0015), liver (HR=5.10, P<0.0001 and HR=2.26, P<0.001), lung (HR=1.54, P=0.011), pancreatic (HR=2.09, P=0.002, HR=1.46, P=0.018, and HR=1.99, P<0.0001) and stomach (HR=1.78, P=0.004) (Fig.2B).

**Figure 2.**
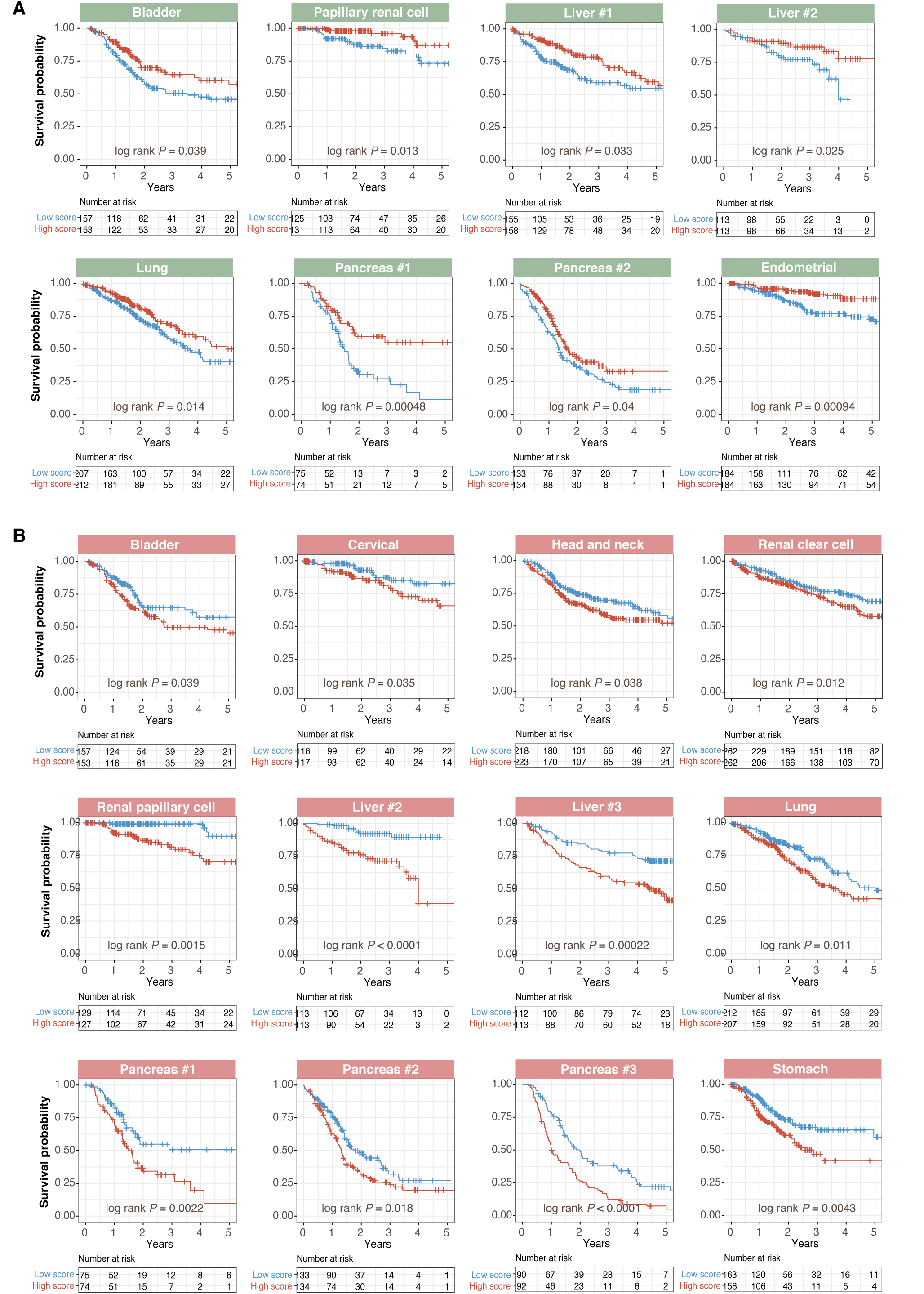
Kaplan-Meier analyses confirming that gene signatures were associated with patients’ overall survival. **(A)** Validation of signature 1 (green panels) across multiple cancer types. Kaplan-Meier plots of overall survival in cancer patients stratified based on signature 1 mean expression scores. Patients were median-dichotomized into high-and low-score groups. Signature 1 is a marker of good prognosis and hence patients with high signature 1 scores had improved survival rates. **(B)** Validation of signature 2 (red panels) across multiple cancer types. Kaplan-Meier plots of overall survival in cancer patients stratified based on signature 2 mean expression scores. Patients were median-dichotomized into high-and low-score groups. Signature 2 is a marker of adverse prognosis and hence patients with high signature 2 scores had poorer survival rates. P-values were calculated from the log-rank test. Pancreas #1 = PAAD cohort; Pancreas #2 = PACA-AU cohort; Pancreas #3 = PACA-CA cohort; Liver #1 = LIHC cohort; Liver #2 = LIRI-JP cohort and Liver #3 = GSE14520 cohort (Table S1).

### Cross-platform subgroup and multivariate analyses confirmed the validity of signatures 1 and 2 as independent prognostic factors

To assess the independence of signatures 1 and 2 over current tumor staging systems, we performed subgroup analyses of their prognostic effect in patients with early (stages 1 and/or 2), intermediate (stages 2 and/or 3) and late (stages 3 and/or 4) cancer stages. Kaplan-Meier analyses revealed that signature 1 successfully identified high-risk (low-expression score) and low-risk (high-expression score) patients with early (bladder, liver, lung, pancreatic and endometrial), intermediate (liver and endometrial) late (papillary renal cell) disease stages (Fig.3A; Fig.S1A). For signature 2, patients with high-expression scores had a significantly poorer survival (Fig.3B). Signature 2 was also independent of disease stage as it successfully predicted survival in early (liver, lung and pancreatic), intermediate (bladder, liver, lung and stomach) and late (bladder, head and neck, papillary renal cell, liver and stomach) stages (Fig.3B; Fig.S1B).

**Figure 3.**
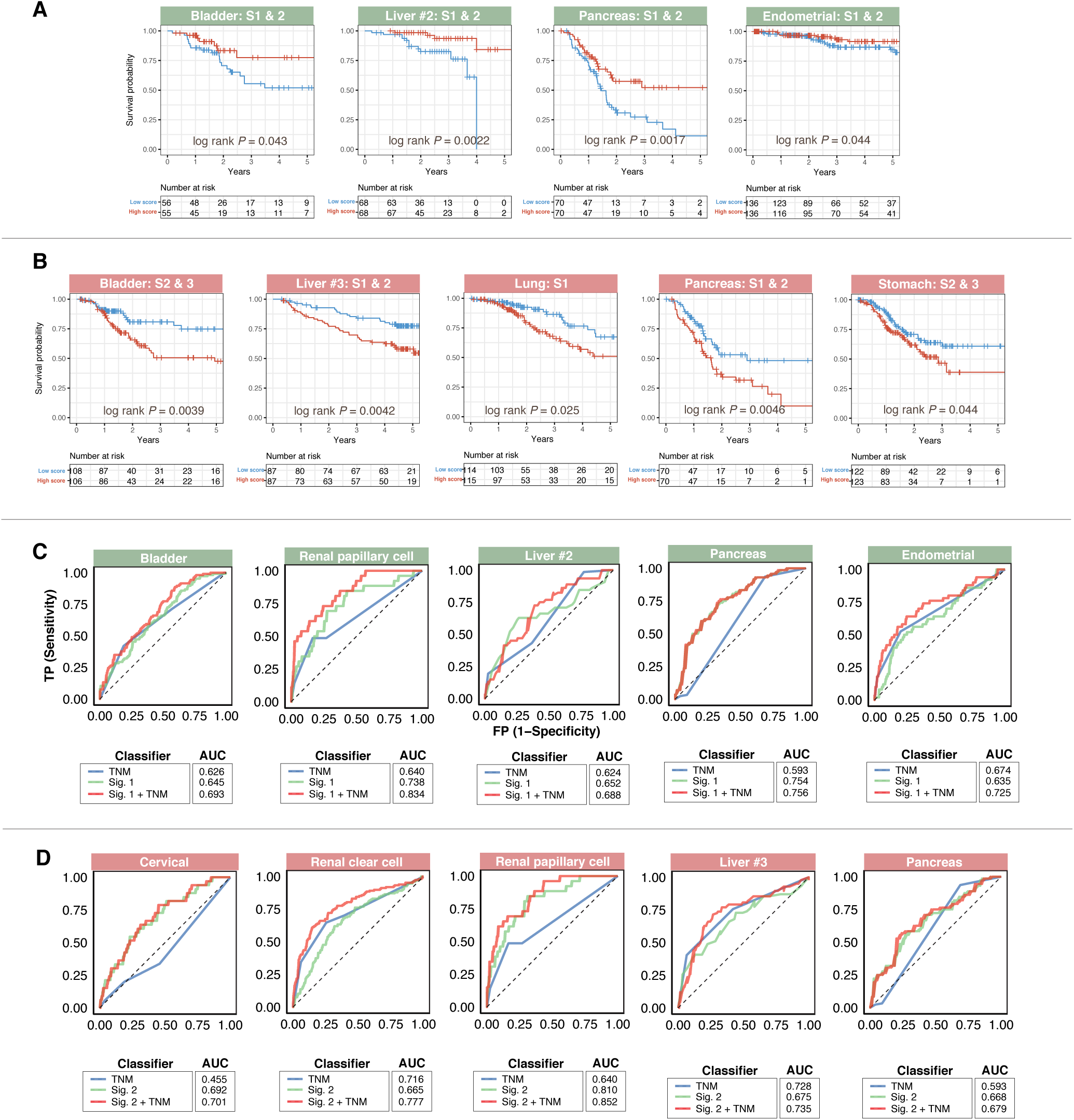
Tumor subgroup analyses and evaluation of predictive performance. Gene signatures were prognostic across different malignant grades. Kaplan-Meier plots show independence of **(A)** signature 1 (green panels) and **(B)** signature 2 (red panels) over current staging systems in cancer cohorts. Patients were sub-grouped according to TNM stages and further stratified using either signature 1 or signature 2 scores. Both signatures successfully identified high-risk patients in different TNM stages. P-values were calculated from the log-rank test. Analysis of specificity and sensitivity of **(C)** signature 1 (green panels) and **(D)** signature 2 (red panels) in cancer cohorts using with Receiver Operating Characteristic (ROC). Plots depict comparison of ROC curves of signatures 1 or 2 and clinical tumor staging parameters. Both signatures demonstrated incremental values over current staging systems. AUC: area under the curve. TNM: tumor, node, metastasis staging. Liver #2 = LIRI-JP cohort and Liver #3 = GSE14520 cohort (Table S1). Representative plots are depicted in this figure. Additional plots are available in Figure S1.

Receiver operating characteristic (ROC) analyses were employed to determine the predictive performance (sensitivity and specificity) of signatures 1 and 2 on 5-year overall survival. Signature 1 performed best, as measured by area under the curve (AUC), in pancreatic cancer (AUC=0.754) followed by papillary renal cell (AUC=0.738), liver (AUC=0.652 and AUC=0.613), bladder (AUC=0.645), endometrial (AUC=0.635) and lung (AUC=0.625) cancers (Fig.3C; Fig.S1C). Signature 2 performance in papillary renal cell cancer (AUC=0.810) was the best, followed by cervical (AUC=0.692), liver (AUC=0.675 and AUC=0.625), pancreatic (AUC=0.668), renal clear cell (AUC=665), head and neck (AUC=0.632), lung (AUC=0.623), stomach (0.618) and bladder (AUC=0.605) cancers (Fig.3D; Fig.S1D). Performance of both signatures 1 and 2 is superior to current tumor staging benchmarks except for the following: signature 1 (liver, lung and endometrial) and signature 2 (bladder, renal clear cell, liver and lung) (Fig.3C,D; Fig.S1C,D). Remarkably, when used in combination with tumor, node and metastasis (TNM) staging parameters, both signatures consistently outperformed each of the individual classifiers (TNM staging or gene signatures when considered alone), reinforcing their incremental prognostic values (Fig.3C,D; Fig.S1C,D). Significantly, while tumor staging could not predict outcome in cervical cancer patients (AUC<0.5), signature 2 sufficiently served as an adverse prognostic factor (Fig.3D).

Univariate Cox regression analyses revealed that tumor stage was associated with survival except for cervical and pancreatic cancers (Table S3). This is expected given the low AUC values for tumor stage in both cancers: pancreatic (AUC=0.593) and cervical (AUC=0.455), suggesting that current staging systems for these cancers are inadequate (Fig.3C,D). Multivariate Cox analyses after adjusting for tumor staging showed that signatures 1 and 2 remained significantly associated with survival (Table S3). For 2 liver cancer cohorts we considered additional clinicopathological features. The GSE14520 cohort consisted of Chinese patients with hepatitis B-associated hepatocellular carcinoma[17] while LIRI-JP is a Japanese-based cohort of mixed etiology[18]. Tumor size, cirrhosis, TNM staging, Barcelona Clinic Liver Cancer staging and alpha-fetoprotein levels were all significantly associated with survival in GSE14520 (Table S3). Tumor size could also predict survival in LIRI-JP (Table S3). When these significant covariates along with signatures 1 or 2 were included in multivariate Cox models, the signatures remained significant risk factors: signature 1 (LIRI-JP: HR=0.54, P=0.043) and signature 2 (LIRI-JP: HR=4.539, P=1.8e-4 and GSE14520: HR=2.01, P=0.003) (Table S3). These results highlight the potentially superior prognostic ability of our signatures; signatures 1 and 2 identified high-and low-risk patients in 8 (n=2,627) and 12 (n=4,134) independent cohorts respectively to collectively represent 10 diverse cancers (Fig.1A).

### Significance of somatic mutations in risk stratified patients

Patients were risk stratified into low-and high-risk groups using signatures 1 and 2. For signature 1, high-risk patients had significantly lower expression levels of good prognosis genes *ALKBH4, ALKBH7, KDM8, KDM6B* and *P4HTM* (Fig.S2A). In contrast, high-risk patients as stratified by signature 2 had significantly higher expression levels of adverse prognosis genes *ASPH, KDM3A, P4HA1, PLOD1* and *PLOD2* (Fig.S2B). To ascertain the relationship between tumor hypoxia and expression of signature genes, hypoxia scores were computed for each patient as mean expression values (log_2_) of 52 hypoxia signature genes[19]. Signature 1 expression scores in patients negatively correlated with hypoxia score (Fig.S3A). Since tumor hypoxia is associated with distant metastasis, recurrence and reduced therapeutic response[20], patients with high expression of signature 1 genes (low hypoxia score) correlated with less advance disease states consistent with it being a marker of good prognosis (Fig.S3A).

Conversely, signature 2 scores positively correlated with tumor hypoxia and hence poor survival outcomes (Fig.S4A). We anticipate that patients’ individual risks of death, as determined from signatures 1 and 2, will positively correlate tumor hypoxia. Indeed, risk scores for each patient, as calculated by taking the sum of Cox regression coefficient for each of the individual genes multiplied with its corresponding expression value[21], significantly correlated with hypoxia scores (Fig.S3B,S4B). Hence, high risk patients have more hypoxic tumors suggesting that our gene signatures are efficient and adequate in predicting death.

To ascertain the association between patients’ risks, as determined by our gene signatures, and somatic mutations, we retrieved the top five most commonly mutated genes for each cancer. Mutations in *PCDHA1*, a cell adhesion gene from the cadherin superfamily, were associated with poor survival in bladder (HR=1.65, P=0.027) and stomach cancers (HR=1.53, P=0.046) but improved survival in endometrial cancer (HR=0.52, P=0.042) (Table S3). Mutations in another gene from the protocadherin alpha cluster, *PCDHA2*, were also associated with adverse outcomes in stomach cancer (HR=1.60, P=0.025) (Table S3). Mutations in *TTN* (connectin) and the tumor suppressor *TP53* were associated with poor survival in bladder (HR=1.61, P=0.016) and endometrial cancers (HR=1.78, P=0.041) (Table S3). Interestingly, another tumor suppressor *PTEN*, when mutated, is linked to better outcomes in endometrial cancer (HR=0.43, P=0.006) (Table S3). Similar observations were made for a lipid kinase gene *PIK3CA* in endometrial cancer (HR=0.36, P=0.002) (Table S3). Likewise, mucin *MUC4* mutations improved survival in clear cell renal cancer patients (HR=0.57, P=0.012) (Table S3); an observation that is consistent with another study[22].

Multivariate Cox analyses on signatures 1 and 2 while controlling for significant somatic mutation variables revealed that the gene signatures were independent survival predictors: bladder (signature 1: HR=0.69, P=0.047 and signature 2: HR=1.58, P=0.043), renal clear cell (signature 2: HR=1.52, P=0.007), stomach (signature 2: HR=1.80, P=0.006) and endometrial (signature 1: HR=0.52, P=0.024) (Table S3). Patients stratified by signatures 1 or 2 and mutation status indicated that they collectively were significantly associated with survival (Fig.4). In bladder cancer, high risk patients (low signature 1 score) harboring mutant alleles of *PCDHA1* had ~50% increased mortality at 5 years compared to low risk patients (high signature 1 score) with wild-type *PCDHA1* (P=0.016) (Fig.4). Although results were less dramatic for *PCDHA1* and signature 2, we still observed a ~25% elevated mortality at 5 years for these two patient groups with bladder cancer (P=0.04) (Fig.4). In patients with stomach cancer, high-risk patients (high signature 2 score) with mutant *PCDHA1* had the worst outcomes (P=0.0017) (Fig.4). Conversely, *PCDHA1* mutation was associated with good prognosis in endometrial cancer, hence high-risk patients with wild-type *PCDHA1* had the lowest survival rates while survival is improved by ~20% in low-risk patients with mutant *PCDHA1* (P=0.003) (Fig.4). *PIK3CA* (P<0.001) and *PTEN* (P=0.0013) mutations were associated with good outcomes in endometrial cancer (Fig.4). Mutations in another cadherin gene *PCDHA2* when considered alongside signature 2 were also associated with survival in stomach cancer (P<0.001) (Fig.4). Survival rates were reduced by ~37% in high-risk patients with mutant *PCDHA2* (Fig.4). Joint relation between *TP53* mutation and signature 1 significantly influenced survival in endometrial cancer (P=0.0015) (Fig.4). Since *MUC4* mutations were associated with better outcomes, survival rates were the lowest in high-risk patients (high signature 2 scores) with wild-type *MUC4* (P=0.0025) (Fig.4).

**Figure 4.**
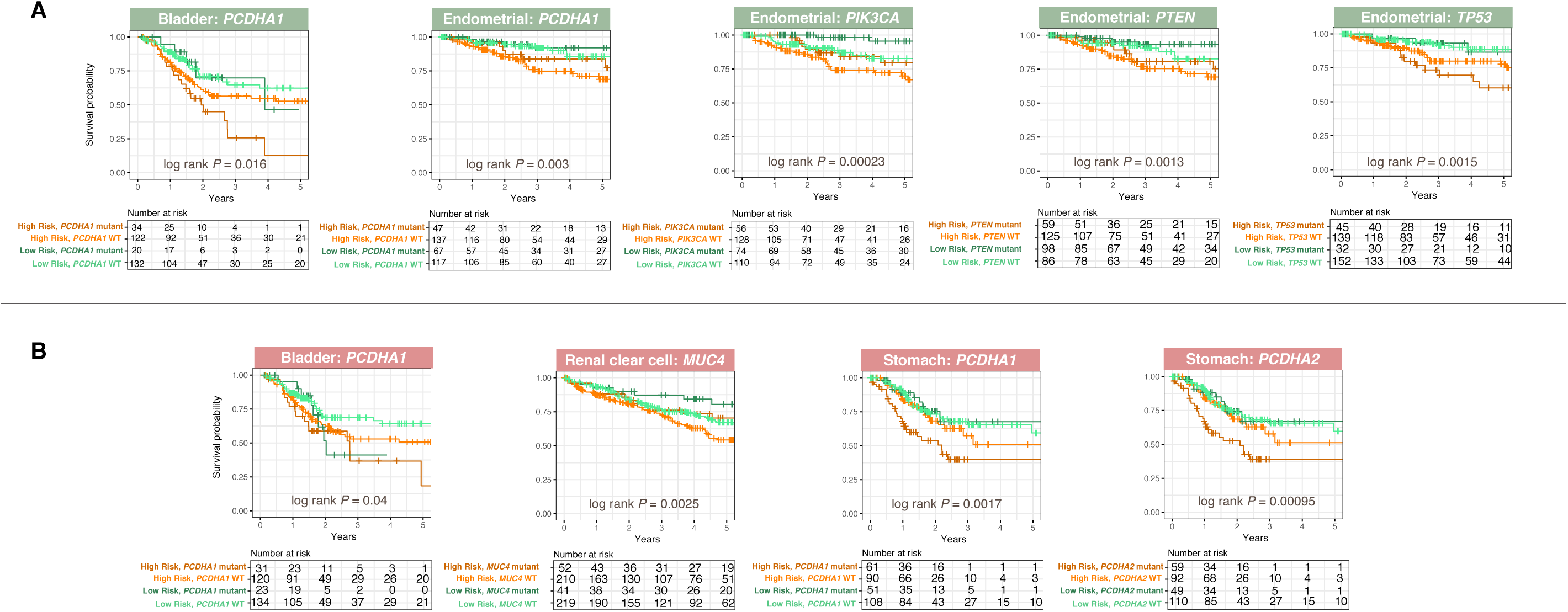
Relationship between patients’ risks as determined by gene signatures and common genetic mutations. Patients were median-stratified into low or high-risk groups using **(A)** signature 1 (green panels) and **(B)** signature 2 (red panels). Since signature 1 is a marker of good prognosis, high-risk patients had lower mean expression of signature 1 genes. Signature 2 is a marker of poor prognosis, hence high-risk patients had higher mean expression of signature 2 genes. Kaplan-Meier plots depicting combined relation of somatic mutations with signatures 1 or 2 on overall survival in cancer patients. P-values were calculated from the log-rank test.

### Tumor suppressive roles of *KDM8* through cell cycle regulation and cell adhesion maintenance

Of all the signature genes, *KDM8* was identified as one of the most downregulated genes in tumors compared to non-tumor samples (Fig.S5A). Patients with high *KDM8* levels had significantly lower risk of death in pancreatic (HR=0.57, P=0.016) and liver cancer cohorts (HR=0.52, P=0.0031; HR=0.49, P=0.026 and HR=0.64, P=0.039) (Fig.5). Prognostic significance of *KDM8* was also independent of tumor stage (Fig.5). *KDM8* expression decreased as tumor malignant grade increased in that the least advanced stage 1 tumors had the highest median *KDM8* values (Fig.S5B). Moreover, *KDM8* expression negatively correlated with hypoxia score, indicating that patients with low levels of *KDM8* had more hypoxic tumors and poorer survival outcomes (Fig.S5C). Together, these observations suggest that *KDM8* may function as a tumor suppressor. This hypothesis is corroborated by an independent report on the role of *KDM8* in cell cycle regulation[23]. Indeed, we observed that *KDM8* expression negatively correlated with the expression levels of canonical cell-cycle genes: cyclins (*CCNA2, CCNB1, CCNB2, CCND1, CCNE1* and *CCNE2*) and cyclin-dependent kinases (*CDK1, CDK2, CDK4, CDK6, CDK7* and *CDK8*), which were consistent across all liver and pancreatic cancer cohorts (Fig.S5D). This implied that *KDM8* is required for tight control of the cell-cycle machinery and its reduction may lead to aberrant proliferation commonly seen in cancer cells.

**Figure 5.**
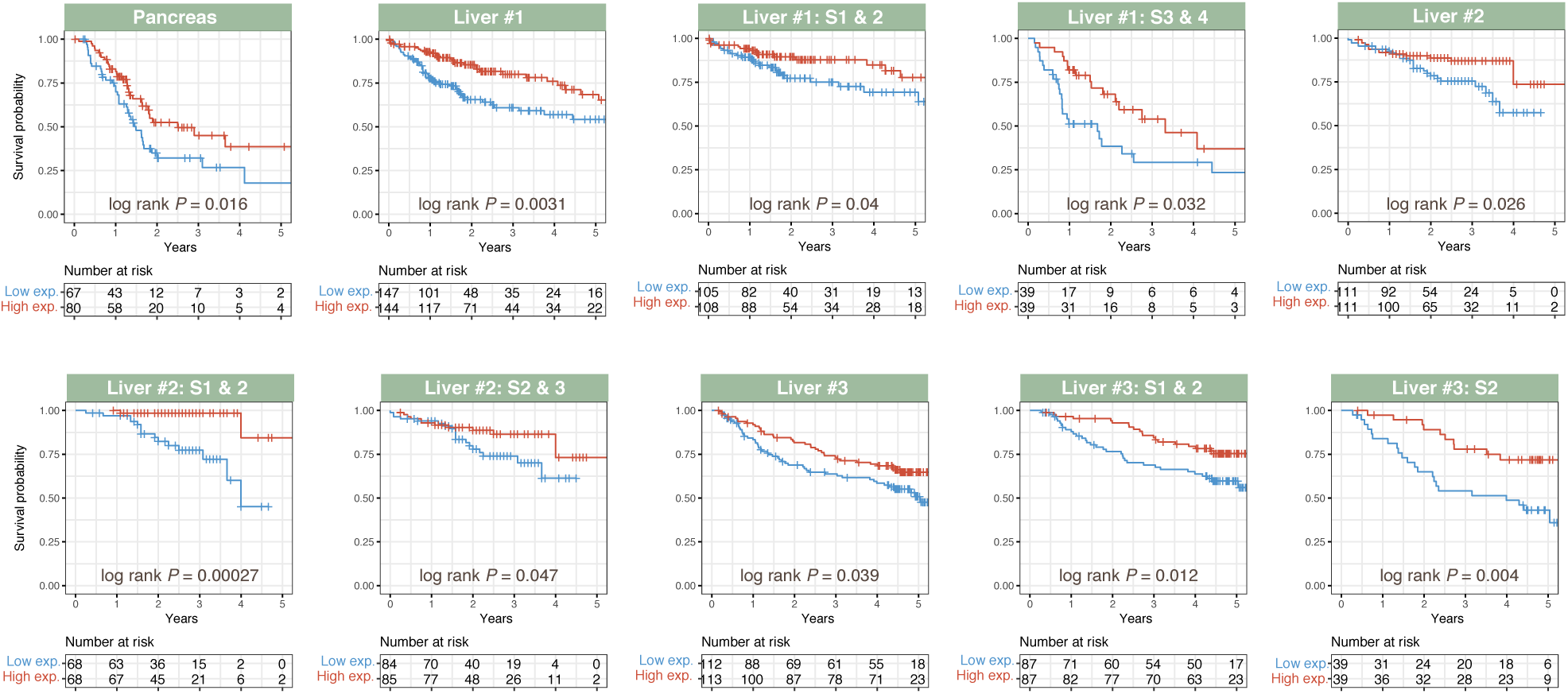
*KDM8* expression is associated with good prognosis in liver and pancreatic cancers. Kaplan-Meier analysis of patients stratified by *KDM8* expression. Patients were median-dichotomized into low-and high-expression groups. Patients with low *KDM8* expression had significantly poorer overall survival. This was consistent in patients analyzed as a full cohort or sub-categorized according to tumor TNM stage. Liver #1 = LIHC cohort; Liver #2 = LIRI-JP cohort and Liver #3 = GSE14520 cohort (Table S1).

To ascertain the biological consequences of deregulated *KDM8* expression, we conducted different expression analysis on liver cancer patients categorized into low and high groups based on *KDM8* expression. A total of 745 genes were differentially expressed (DEGs) between the two groups (fold change>2 or <-2, P<0.05) (Table S4). Significant enrichments of biological pathways involved in metabolism, immune regulation, *VEGF* production, inflammation and cell adhesion were observed (Fig.S5E;Table S5). Furthermore, DEGs were overrepresented as targets of *HNF4A, HNF4G, FOXA1, FOXA2* and *NR2F2* transcription factors (TFs) (Fig.S5F). These TFs play central roles in cell polarity maintenance and epithelial differentiation[24–26], hence downregulation of *KDM8* may drive epithelial-mesenchymal transition (EMT) and tumor progression. *HNF4A* is one of the key TFs responsible for regulating a myriad of hepatic functions including cell junction assembly[26,27]. Pathway analysis of *KDM8* DEGs revealed enrichment of processes related to cell adhesion, suggesting a potential crosstalk between *KDM8* and *HNF4A*. Of the 745 DEGs, analysis on a hepatoma-based HNF4A chromatin immunoprecipitation-sequencing dataset demonstrated that 148 genes were directly bound by HNF4A[28]. To further reinforce the interplay between *HNF4A* and *KDM8*, we observed that 110 of the 745 DEGs were overrepresented in HNF4A-null mice[26] and 45 of these genes were direct HNF4A targets (Fig.S5G). Differential expression analysis between *HNF4A*-deficient and wild-type mice showed that a majority of the 45 genes were downregulated, as expected, suggesting that HNF4A directly activates their gene expression, many of which are involved in a multitude of cell adhesion processes (Fig.S5H).

## Discussion

A multi-cohort retrospective study led to the identification of two novel pan-cancer prognostic gene signatures derived from oxygen-sensing genes. Cross-platform validations confirmed prognosis in 10 cancers to collectively include 6,761 patients spanning 20 diverse cohorts (Fig.1A). The gene signatures had opposing prognostic value; signature 1 is a marker of good prognosis while signature 2 is associated with poor outcomes. The key strengths of our signatures as powerful prognostic tools are: 1) pan-cancer utility, 2) involving a mere 5 genes each that provide continuous assessment of death risks and 3) superior to current tumor staging parameters. The prognostic capability of our signatures in diverse cancers is evidence that conserved biological processes; oxygen-sensing, cell-cycle deregulation and loss of cell polarity, governs disease progression and thus clinical outcomes.

Anti-tumorigenic functions have been reported for several genes from signature 1. Loss of *KDM6B* resulted in more aggressive pancreatic ductal adenocarcinoma[29]. In colorectal cancer, high *KDM6B* is a predictor of good prognosis and knock-down of *KDM6B* augmented cell proliferation and inhibited apoptosis[30]. Yet, *KDM6B* function is enigmatic. Others have reported that high *KDM6B* expression is associated with increased metastasis and invasion in renal clear cell carcinoma that correlated with poor prognosis[31]. *KDM6B* also promotes TGF-B induced EMT and invasiveness in breast cancer[32]. While we could neither confirm nor deny the validity of these studies, it is striking that our observation of favorable prognosis associated with high *KDM6B* in pancreatic ductal adenocarcinoma was consistent with the report from Yamamoto and colleagues[29]. We also did not observe any prognostic significance of signature 1, in which *KDM6B* is part of, in either breast or renal clear cell cancers, which indirectly substantiates findings from the two other reports on *KDM6B* not being a marker of good prognosis[31,32].

*KDM8* is a gene associated with favorable prognosis. Our results suggest determinative crosstalk between *KDM8* and *HNF4A*, particularly in the context of morphogenesis, cell adhesion, maintenance of cell polarity and epithelial formation. Moreover, survival rates for bladder cancer at 5 years dropped to ~12% in patients that simultaneously harbor low expression of signature 1 genes (high-risk), which included *KDM8*, and *PCDHA1* mutations. Additive effects conferred by mutations in this cell-adhesion protein provides support for the hypothesis that *KDM8* is likely a tumor suppressor and downregulation of this gene may lead to a loss of epithelial phenotype and cell adhesion to promote cancer invasion. Additionally, loss of *KDM8* expression is correlated with increased expression of cell-cycle genes that may contribute to deranged cell cycle regulation and tumor progression (Fig.S5D). Like *KDM6B*, the function of *KDM8* appears to be cell-type dependent. While *KDM8* expression is downregulated in liver and pancreatic tumors compared to adjacent cells (Fig.S5A), it is overexpressed in breast cancer to induce EMT and invasion[33]. Nonetheless, *KDM8* roles are not limited to cell cycle regulation. *KDM8* exert tumor suppressive functions in hematopoietic cancer by mediating DNA repair[34]. Collectively, imbalance in the JmjC subfamily of lysine demethylases such as *KDM8* and *KDM6B* are likely to result in broad-ranging but cell-type specific biological effects.

Overall, our gene signatures will enhance decision making in clinic and provide individualized treatment plans based on patients’ tumor biology to maximize treatment efficacy and prolong lifespan by directing resources to those most in need. Upon diagnosis, the signatures can be used for risk stratification and patients with poor prognosis can be subjected to additional neoadjuvant treatments. These signatures would need to be assessed prospectively and we aim to evaluate their use in clinical decision making of patients presented with these cancers.

## Acknowledgements

A special thank you to Graham Foster for his critical insights and help on manuscript preparation.

## Authors contribution

WHC and AGL designed the study and analyzed the data. All authors interpreted the data. WHC and AGL wrote the initial manuscript draft. All authors revised the manuscript and approved the final version.

## Supplementary methods

### Study cohorts

Cancer datasets used in this study were obtained from The Cancer Genome Atlas (TCGA)[15], International Cancer Genome Consortium[16] and Gene Expression Omnibus (n=20,752). Cohort descriptions were listed in Table S1. Gene expression profiles for TCGA cancers were downloaded from Broad Institute GDAC Firehose. Gene expression profiles for ICGC cancers were downloaded from the ICGC data portal. TCGA Illumina HiSeq rnaseqv2 Level 3 RSEM normalized gene expression and ICGC normalized gene expression profiles were converted to log_2_(x + 1) scale.

### *KDM8* differential expression analysis

TCGA liver cancer cohort (LIHC; Table S1) was used. Patients (n=371) were median dichotomized into low and high groups based on *KDM8* expression. To determine differentially expressed genes between both groups, the Bayes method and linear model were employed using the R package limma. P-values were adjusted using the false discovery rate controlling procedure of Benjamini-Hochberg. Genes with log_2_ fold change of > 1 or < -1 and adjusted P-values < 0.05 were considered significant.

### Gene signature and risk scores

Expression scores for signature 1 and 2 were calculated for each patient by taking the average log_2_ expression values of signature genes. As a measure of tumor hypoxia, hypoxia scores were determined by calculating the average log_2_ expression values of 52 hypoxia signature genes[19]. Risk scores for each patient were determined by taking the sum of Cox regression coefficient for each of the signature genes multiplied with its corresponding expression value. Nonparametric Spearman’s rank correlation was employed to assess the relationship of expression scores and risk scores with tumor hypoxia (hypoxia score).

### Survival analyses

Cox proportional hazards regression was employed to investigate the association between patient survival duration and one or more risk factors, e.g. signature 1 or signature 2 scores, tumor stage and other clinical variables. Univariate analyses were performed to determine the influence of individual risk factors on overall survival. Multivariate analyses were performed by including risk factors that were significantly associated with overall survival identified in univariate analyses (P < 0.05) since multiple covariates can influence patient prognosis. Hazard ratios (HR) were determined from Cox models with HR greater than one indicating that a covariate was positively associated with event probability (increased hazard) and negatively associated with survival length. Cox regression analyses were performed using the R survival and survminer packages. Proportional hazards assumption was supported by a non-significant relationship between scaled Schoenfeld residuals and time using the R survival package. In addition, Kaplan-Meier and log-rank tests were used in univariate analyses of the gene signatures in relation to patient survival and were performed using the survival and survminer packages. Patients were median-dichotomized into low and high-score groups based on mean expression scores of signature genes. Difference between high and low-score groups were tested using the log-rank test implemented with the survival package.

Time-dependent receiver operating characteristic (ROC) curve analysis was used to assess the predictive performance of both signatures 1 and 2 in comparison with standard tumor staging parameters. The R survcomp package was employed to compute time-dependent ROC curves[35]. ROC curves depicted true positive rates (sensitivity) versus false positive rates (1-specificity). The area under the curve (AUC) is a measure of how well the gene signatures can predict patient survival where AUC ranges from 0.5 (for an uninformative marker) to 1 (for a perfect predictive marker).

### Biological enrichment analysis

Analysis of biological pathway enrichment on the 745 differentially expressed genes between *KDM8*-low and-high groups was conducted using GeneCodis against the Kyoto Encyclopedia of Genes and Genomes (KEGG) and Gene Ontology (GO) databases[36]. The Enrichr tool was used to identify transcription factors from the ENCODE database that are potential regulators of these 745 genes[37,38].

### *HNF4A* loss of function analysis

A total of 148 genes were identified as *HNF4A* targets in the HepG2 hepatoma cell line determined using the Enrichr tool[37,38]. Differential expression analysis between *HNF4A* wild-type and null mice livers (GSE3126) performed using the GEO2R tool[26] identified 110 differentially expressed genes from the initial 745-gene set identified previously (Fig.S5G). Of these 110 genes, 45 genes were identified as direct *HNF4A* targets and were downregulated in the *HNF4A*-null mice (Fig.S5H).

### Somatic mutation identification

Level 3 mutation datasets were downloaded from GDAC. Kaplan-Meier analysis and log-rank tests were employed to determine the association of somatic mutations in combination with the signatures 1 or 2, on overall survival.

All graphs were generated using the ggplot2 package in R[39].

**Figure S1.**
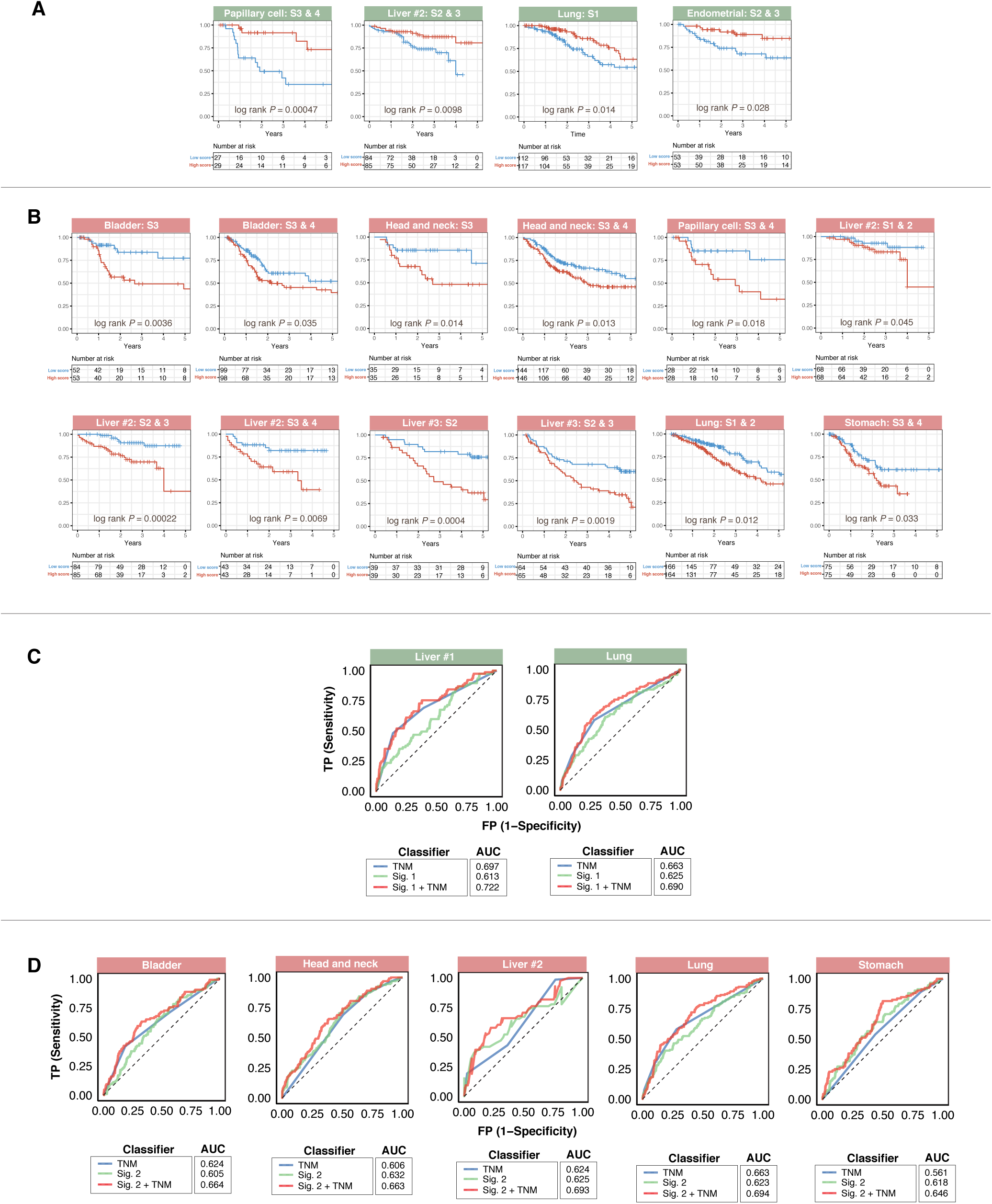
Additional tumor subgroup analyses and evaluation of predictive performance. Additional plots from Figure 3. Gene signatures were prognostic across different malignant grades. Kaplan-Meier plots show independence of **(A)** signature 1 (green panels) and **(B)** signature 2 (red panels) over current staging systems in cancer cohorts. Patients were sub-grouped according to TNM stages and further stratified using either signature 1 or signature 2 scores. Both signatures successfully identified high-risk patients in different TNM stages. P-values were calculated from the log-rank test. Analysis of specificity and sensitivity of **(C)** signature 1 (green panels) and **(D)** signature 2 (red panels) in cancer cohorts using with Receiver Operating Characteristic (ROC). Plots depict comparison of ROC curves of signatures 1 or 2 and clinical tumor staging parameters. Both signatures demonstrated incremental values over current staging systems. AUC: area under the curve. TNM: tumor, node, metastasis staging. Liver #1 = LIHC cohort; Liver #2 = LIRI-JP cohort and Liver #3 = GSE14520 cohort (Table S1).

**Figure S2.**
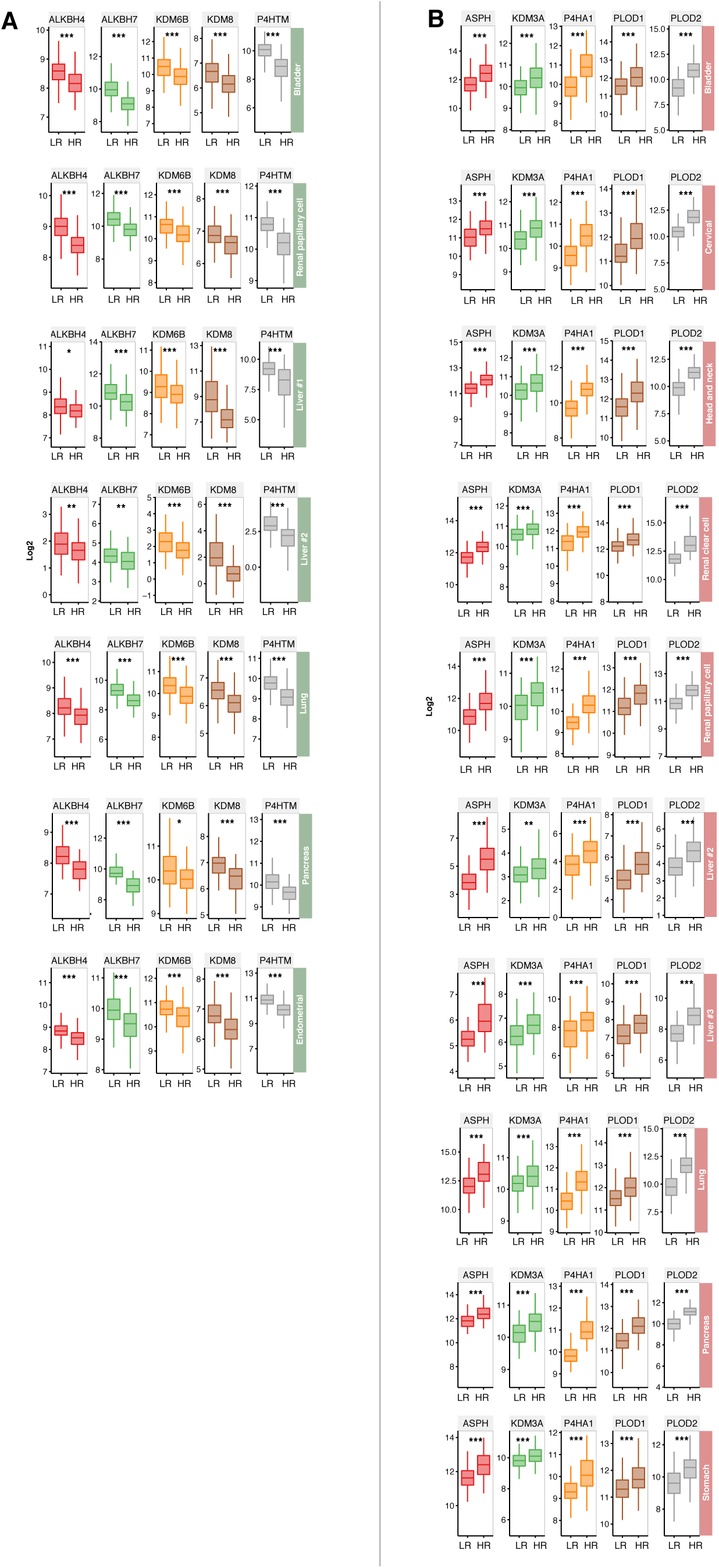
Expression of distribution of signature genes in low-and high-risk patients. **(A)** signature 1 (green panels) and **(B)** signature 2 (red panels). Patients were median-stratified into low-and high-risk groups based on mean expression of signature genes. Box plots depict expression distribution of each of the 5 genes in both signatures in these two patient groups. **(A)** Since signature 1 is a marker of good prognosis, high-risk patients showed significantly lower expression of individual signature genes. **(B)** In contrast, signature 2 is a marker of poor prognosis, hence high-risk patients showed significantly higher expression of individual signature genes. Nonparametric Mann-Whitney-Wilcoxon tests were used to compare low-and high-risk patients. Asterisks represent significant P values: * < 0.01, ** < 0.001 and *** < 0.0001. LR = low risk. HR = high risk.

**Figure S3.**
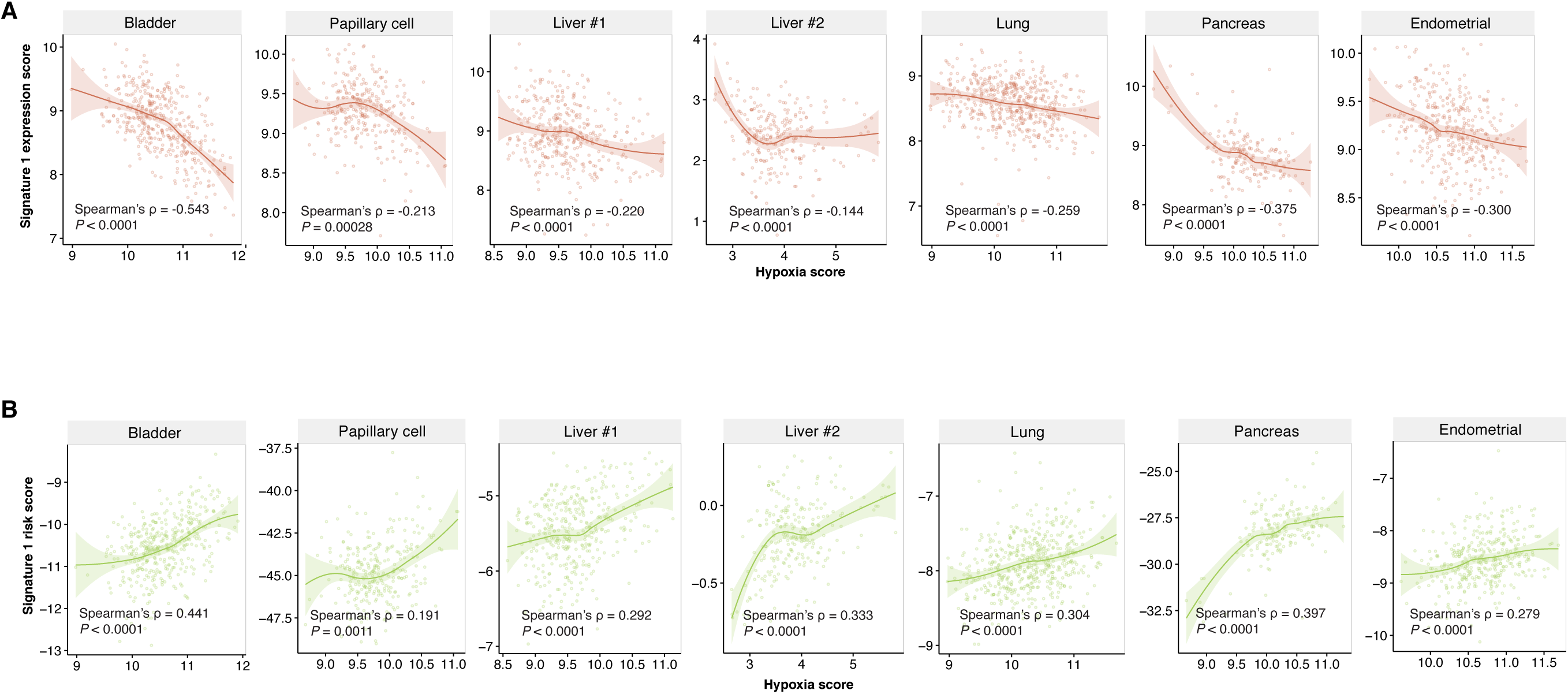
Correlation of patients’ risk scores derived from signature 1 with tumor hypoxia. **(A)** Significant negative correlation between signature 1 expression scores and tumor hypoxia. **(B)** Significant positive correlation between signature 1 risk scores and tumor hypoxia. Calculations of expression scores, risk scores and hypoxia scores are explained in the methods. Liver #1 = LIHC cohort and Liver #2 = LIRI-JP cohort.

**Figure S4.**
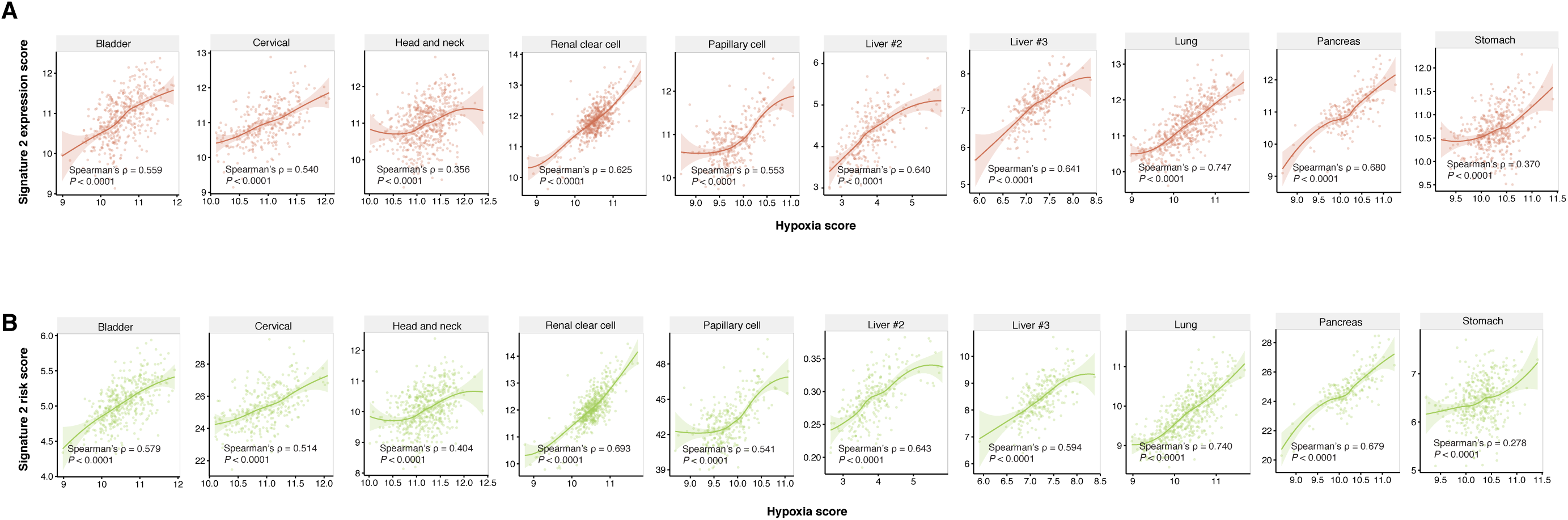
Correlation of patients’ risk scores derived from signature 2 with tumor hypoxia. (A) Significant positive correlation between signature 2 expression scores and tumor hypoxia. **(B)** Significant positive correlation between signature 2 risk scores and tumor hypoxia. Calculations of expression scores, risk scores and hypoxia scores are explained in the methods. Liver #2 = LIRI-JP cohort and Liver #3 = GSE14520 cohort.

**Figure S5.**
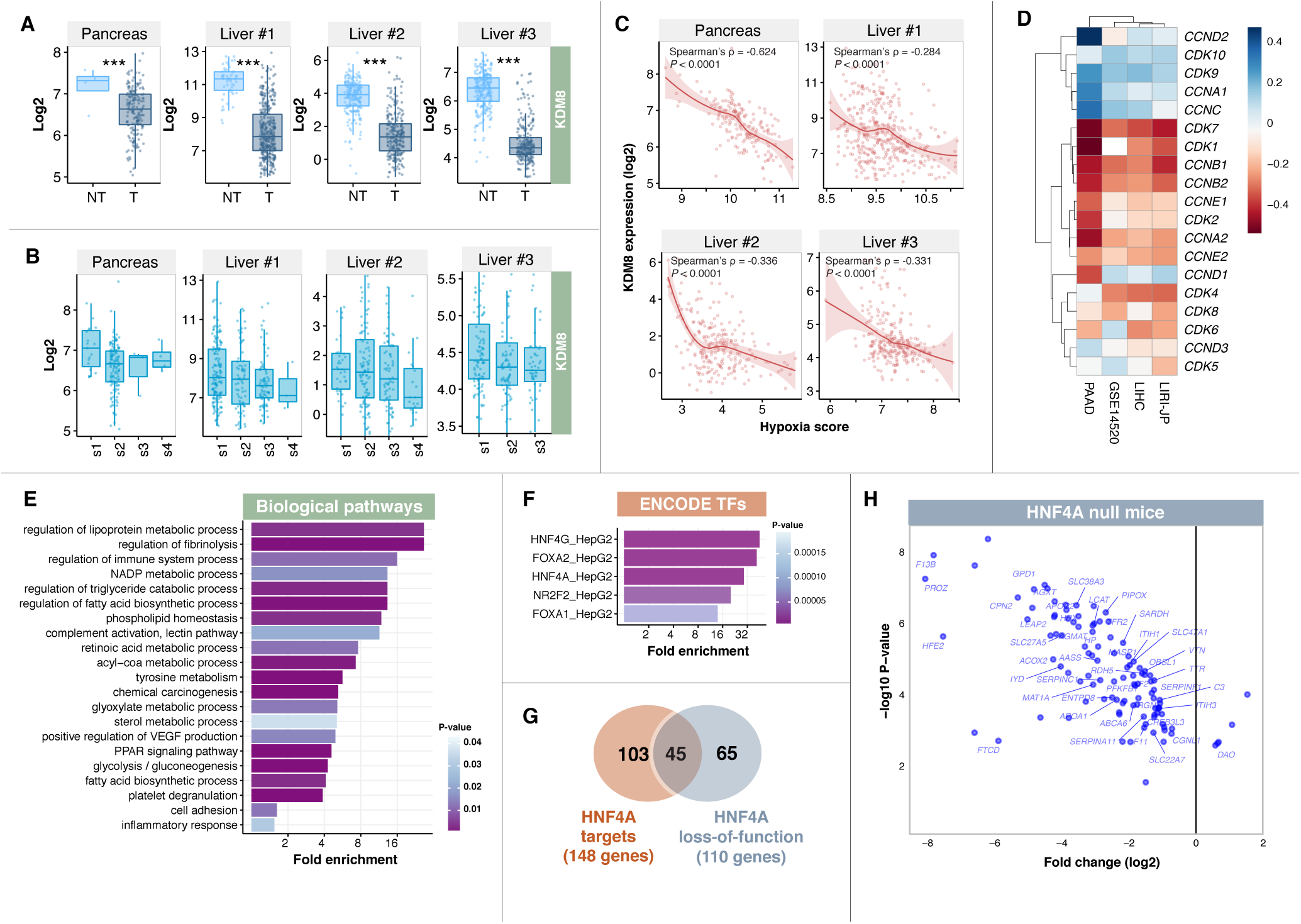
Putative tumor suppressive functions of *KDM8* occur through processes related to cell cycle regulation and cell adhesion. **(A)** Expression distributions of *KDM8* were significantly lower in tumor (T) compared to non-tumor (NT) samples in liver and pancreatic cancer cohorts. Mann-Whitney-Wilcoxon tests were used to compare T and NT samples. Asterisks represent significant P values: *** < 0.0001. **(B)** Expression levels of *KDM8* decreased according to disease progression and malignant grade in liver and pancreatic cancer cohorts. **(C)** Significant negative correlation between patients’ *KDM8* expression and tumor hypoxia (hypoxia score) in liver and pancreatic cancer cohorts. **(D)** Correlation between *KDM8* expression and canonical cell-cycle regulators in patients with liver or pancreatic cancers. A majority of genes involved in cell-cycle regulation are negatively correlated with *KDM8* expression. Liver #1 = LIHC cohort; Liver #2 = LIRI-JP cohort and Liver #3 = GSE14520 cohort (Table S1). **(E)** Patients were median-stratified according to KDM8 expression. Differential expression analysis between *KDM8* high-and low-groups in liver cancer revealed 745 differentially expressed genes (DEGs; fold-change >2 or <-2). Enrichment of biological pathways associated with DEGs, which include processes related to cell adhesion, inflammation, metabolism and signal transduction pathways related to cancer. **(F)** Enrichment of transcription factors (TFs) from the ENCODE database that are potential regulators of *KDM8* DEGs. These TFs were predicted to bind near *KDM8* DEGs. **(G)** Venn diagram depicts the overlap between HNF4A targets (as identified by ENCODE chromatin-immunoprecipitation sequencing dataset) and genes affected by HNF4A loss-of-function (as identified in HNF4A-null mice). Of the 745 DE genes, 148 genes were identified as direct HNF4A targets and 110 genes were affected by HNF4A loss-of-function. In the Venn intersection, 45 genes were both HNF4A targets and altered in HNF4A-null mice. **(H)** Scatter plot depicts expression patterns of 110 genes affected by HNF4A loss-of-function. Gene names of the 45 HNF4A targets were annotated on the plot. A majority of *KDM8*-associated genes were downregulated in the HNF4A-null mice.

**Table S1.** Cancer cohort descriptions.

| **Cohort** | **Non-tumour #** | **Tumour #** | **Description** |
| --- | --- | --- | --- |
| BLCA | 19 | 408 | Bladder Urothelial Carcinoma |
| BRCA | 112 | 10939 | Breast invasive carcinoma |
| CESC | 3 | 304 | Cervical squamous cell carcinoma and endocervical adenocarcinoma |
| CHOL | 9 | 36 | Cholangiocarcinoma |
| COAD | 41 | 285 | Colon adenocarcinoma |
| ESCA | 11 | 184 | Esophageal carcinoma |
| GBM | 5 | 153 | Glioblastoma multiforme |
| GBMLGG | 5 | 669 | Glioma |
| HNSC | 44 | 520 | Head and Neck squamous cell carcinoma |
| KICH | 25 | 66 | Kidney Chromophobe |
| KIPAN | 129 | 889 | Pan-kidney cohort |
| KIRC | 72 | 533 | Kidney renal clear cell carcinoma |
| KIRP | 32 | 290 | Kidney renal papillary cell carcinoma |
| LIHC (Liver cohort 1) | 50 | 371 | Hepatocellular carcinoma |
| LUAD | 59 | 515 | Lung adenocarcinoma |
| LUSC | 51 | 501 | Lung squamous cell carcinoma |
| PAAD (Pancreas cohort 1) | 4 | 178 | Pancreatic adenocarcinoma |
| PCPG | 3 | 179 | Pheochromocytoma and Paraganglioma |
| PRAD | 52 | 497 | Prostate adenocarcinoma |
| SARC | 2 | 259 | Sarcoma |
| STAD | 35 | 415 | Stomach adenocarcinoma |
| STES | 46 | 599 | Stomach and Esophageal carcinoma |
| THCA | 59 | 501 | Thyroid carcinoma |
| THYM | 2 | 120 | Thymoma |
| UCEC | 11 | 370 | Uterine Corpus Endometrial Carcinoma |
| LIRI-JP (Liver cohort 2) | NA | 226 | Hepatocellular carcinoma |
| GSE14520 (Liver cohort 3) | NA | 242 | Hepatocellular carcinoma |
| PACA-AU (Pancreas cohort 2) | NA | 269 | Pancreatic adenocarcinoma |
| PACA-CA (Pancreas cohort 3) | NA | 234 | Pancreatic adenocarcinoma |

**Table S2.** List of 61 2OG oxygenases.

| Gene Symbol | Description |
| --- | --- |
| ALKBH1 | alkB homolog 1, histone H2A dioxygenase |
| ALKBH2 | alkB homolog 2, alpha-ketoglutarate dependent dioxygenase |
| ALKBH3 | alkB homolog 3, alpha-ketoglutaratedependent dioxygenase |
| ALKBH4 | alkB homolog 4, lysine demethylase |
| ALKBH5 | alkB homolog 5, RNA demethylase |
| ALKBH6 | alkB homolog 6 |
| ALKBH7 | alkB homolog 7 |
| ALKBH8 | alkB homolog 8, tRNA methyltransferase |
| ASPH | aspartate beta-hydroxylase |
| BBOX1 | gamma-butyrobetaine hydroxylase 1 |
| EGLN1 | egl-9 family hypoxia inducible factor 1 |
| EGLN2 | egl-9 family hypoxia inducible factor 2 |
| EGLN3 | egl-9 family hypoxia inducible factor 3 |
| FTO | FTO, alpha-ketoglutarate dependent dioxygenase |
| HR | HR, lysine demethylase and nuclear receptor corepressor |
| HSPBAP1 | HSPB1 associated protein 1 |
| JARID2 | jumonji and AT-rich interaction domain containing 2 |
| JMJD1C | jumonji domain containing 1C |
| JMJD4 | jumonji domain containing 4 |
| JMJD6 | arginine demethylase and lysine hydroxylase |
| JMJD7_PLA2G4B | jumonji Domain Containing 7-Phospholipase A2, Group IVB (Cytosolic) Read-Through |
| JMJD8 | jumonji domain containing 8 |
| KDM1A | lysine demethylase 1A |
| KDM1B | lysine demethylase 1B |
| KDM2A | lysine demethylase 2A |
| KDM2B | lysine demethylase 2B |
| KDM3A | lysine demethylase 3A |
| KDM3B | lysine demethylase 3B |
| KDM4A | lysine demethylase 4A |
| KDM4B | lysine demethylase 4B |
| KDM4C | lysine demethylase 4C |
| KDM4D | lysine demethylase 4D |
| KDM4E | lysine demethylase 4E |
| KDM5A | lysine demethylase 5A |
| KDM5B | lysine demethylase 5B |
| KDM5C | lysine demethylase 5C |
| KDM5D | lysine demethylase 5D |
| KDM6A | lysine demethylase 6A |
| KDM6B | lysine demethylase 6B |
| KDM7A | lysine demethylase 7A |
| KDM8 | lysine demethylase 8 |
| OGFOD1 | 2-oxoglutarate and iron dependent oxygenase domain containing 1 |
| P3H1 | prolyl 3-hydroxylase 1 |
| P3H2 | prolyl 3-hydroxylase 2 |
| P3H3 | prolyl 3-hydroxylase 3 |
| P4HA1 | prolyl 4-hydroxylase subunit alpha 1 |
| P4HA2 | prolyl 4-hydroxylase subunit alpha 2 |
| P4HA3 | prolyl 4-hydroxylase subunit alpha 3 |
| P4HTM | prolyl 4-hydroxylase, transmembrane |
| PHF2 | PHD finger protein 2 |
| PHF8 | PHD finger protein 8 |
| PHYH | phytanoyl-CoA 2-hydroxylase |
| PHYHD1 | phytanoyl-CoA dioxygenase domain containing 1 |
| PLOD1 | procollagen-lysine,2-oxoglutarate 5-dioxygenase 1 |
| PLOD2 | procollagen-lysine,2-oxoglutarate 5-dioxygenase 2 |
| PLOD3 | procollagen-lysine,2-oxoglutarate 5-dioxygenase 3 |
| TET1 | tet methylcytosine dioxygenase 1 |
| TET2 | tet methylcytosine dioxygenase 2 |
| TET3 | tet methylcytosine dioxygenase 3 |
| TMLHE | trimethyllysine hydroxylase, epsilon |
| UTY | ubiquitously transcribed tetratricopeptide repeat containing, Y-linked |

**Table S3.**
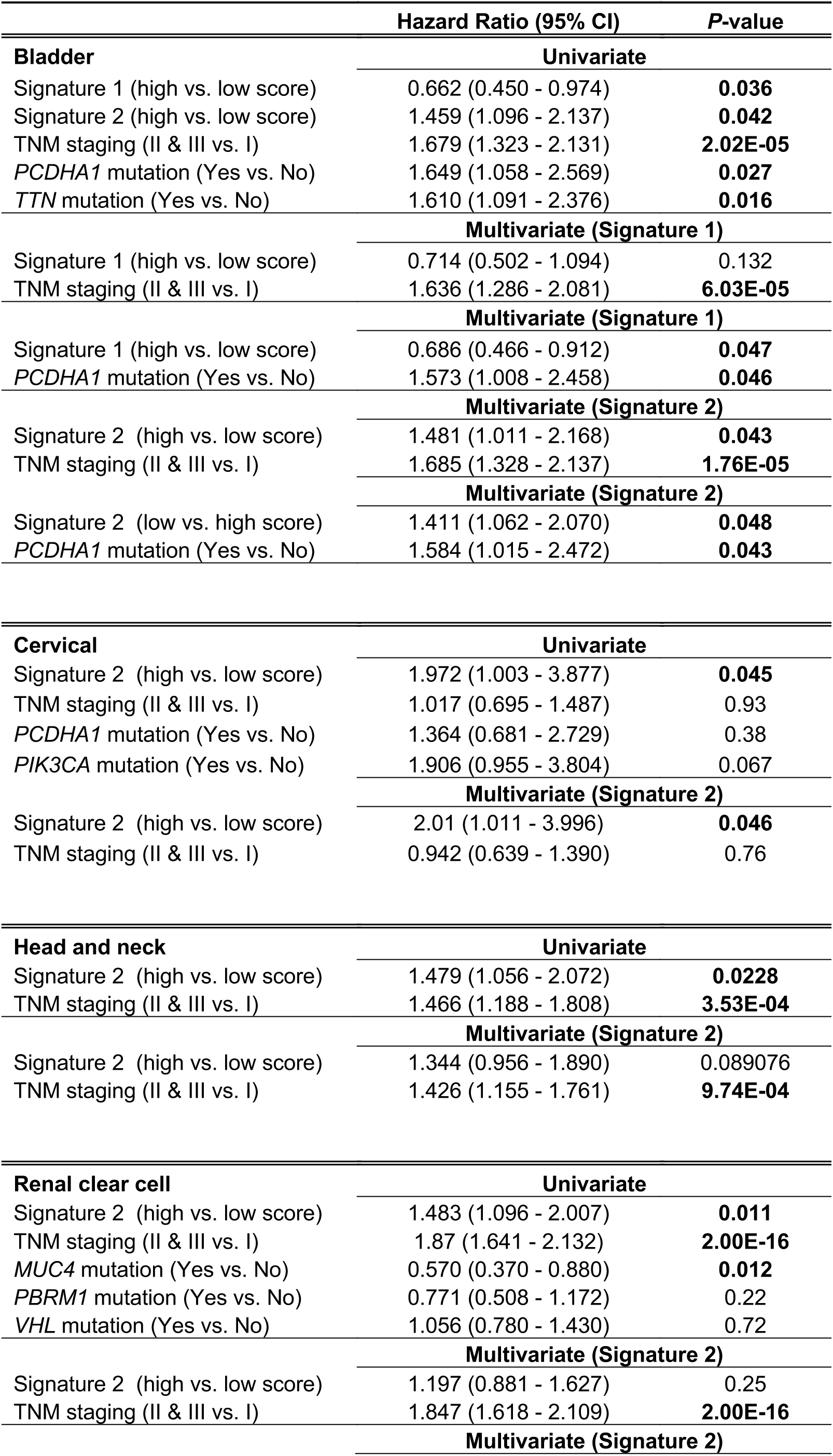
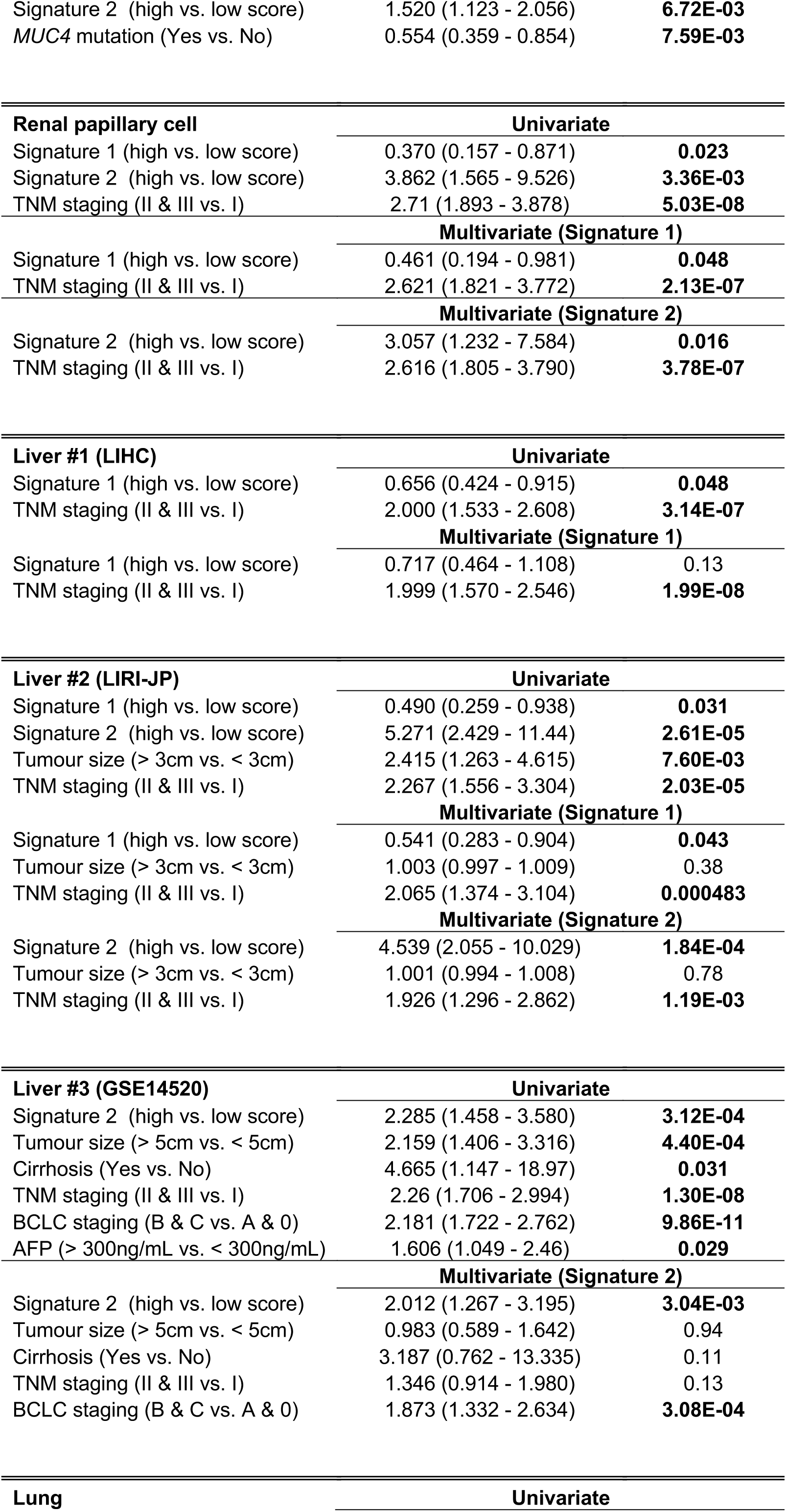
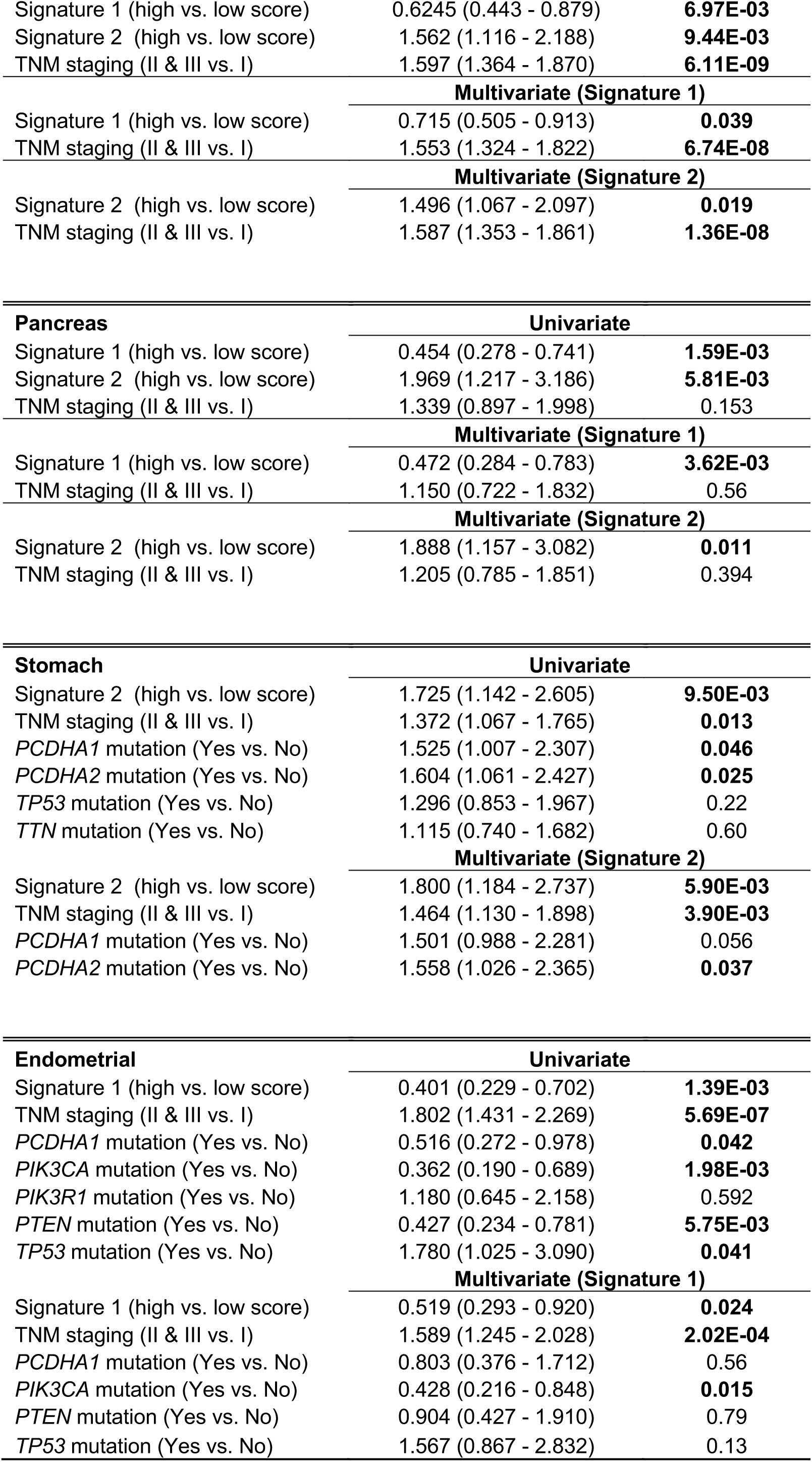
Univariate and multivariate Cox proportional hazards analysis of risk factors associated with overall survival in multiple cancers. Refer to Table S1 for cancer abbreviations.

|  | Hazard Ratio (95% CI) | P-value |
| --- | --- | --- |
| **Bladder** | **Univariate** |  |
| Signature 1 (high vs. low score) | 0.662 (0.450 - 0.974) | **0.036** |
| Signature 2 (high vs. low score) | 1.459 (1.096 - 2.137) | **0.042** |
| TNM staging (II & III vs. I) | 1.679 (1.323 - 2.131) | **2.02E-05** |
| *PCDHA1* mutation (Yes vs. No) | 1.649 (1.058 - 2.569) | **0.027** |
| *TTN* mutation (Yes vs. No) | 1.610 (1.091 - 2.376) | **0.016** |
|  | **Multivariate (Signature 1)** |  |
| Signature 1 (high vs. low score) | 0.714 (0.502 - 1.094) | 0.132 |
| TNM staging (II & III vs. I) | 1.636 (1.286 - 2.081) | **6.03E-05** |
|  | **Multivariate (Signature 1)** |  |
| Signature 1 (high vs. low score) | 0.686 (0.466 - 0.912) | **0.047** |
| *PCDHA1* mutation (Yes vs. No) | 1.573 (1.008 - 2.458) | **0.046** |
|  | **Multivariate (Signature 2)** |  |
| Signature 2 (high vs. low score) | 1.481 (1.011 - 2.168) | **0.043** |
| TNM staging (II & III vs. I) | 1.685 (1.328 - 2.137) | **1.76E-05** |
|  | **Multivariate (Signature 2)** |  |
| Signature 2 (low vs. high score) | 1.411 (1.062 - 2.070) | **0.048** |
| *PCDHA1* mutation (Yes vs. No) | 1.584 (1.015 - 2.472) | **0.043** |
| **Cervical** | **Univariate** |  |
| Signature 2 (high vs. low score) | 1.972 (1.003 - 3.877) | **0.045** |
| TNM staging (II & III vs. I) | 1.017 (0.695 - 1.487) | 0.93 |
| *PCDHA1* mutation (Yes vs. No) | 1.364 (0.681 - 2.729) | 0.38 |
| *PIK3CA* mutation (Yes vs. No) | 1.906 (0.955 - 3.804) | 0.067 |
|  | **Multivariate (Signature 2)** |  |
| Signature 2 (high vs. low score) | 2.01 (1.011 - 3.996) | **0.046** |
| TNM staging (II & III vs. I) | 0.942 (0.639 - 1.390) | 0.76 |
| **Head and neck** | **Univariate** |  |
| Signature 2 (high vs. low score) | 1.479 (1.056 - 2.072) | **0.0228** |
| TNM staging (II & III vs. I) | 1.466 (1.188 - 1.808) | **3.53E-04** |
|  | **Multivariate (Signature 2)** |  |
| Signature 2 (high vs. low score) | 1.344 (0.956 - 1.890) | 0.089076 |
| TNM staging (II & III vs. I) | 1.426 (1.155 - 1.761) | **9.74E-04** |
| **Renal clear cell** | **Univariate** |  |
| Signature 2 (high vs. low score) | 1.483 (1.096 - 2.007) | **0.011** |
| TNM staging (II & III vs. I) | 1.87 (1.641 - 2.132) | **2.00E-16** |
| *MUC4* mutation (Yes vs. No) | 0.570 (0.370 - 0.880) | **0.012** |
| *PBRM1* mutation (Yes vs. No) | 0.771 (0.508 - 1.172) | 0.22 |
| *VHL* mutation (Yes vs. No) | 1.056 (0.780 - 1.430) | 0.72 |
|  | **Multivariate (Signature 2)** |  |
| Signature 2 (high vs. low score) | 1.197 (0.881 - 1.627) | 0.25 |
| TNM staging (II & III vs. I) | 1.847 (1.618 - 2.109) | **2.00E-16** |
|  | **Multivariate (Signature 2)** |  |
| Signature 2 (high vs. low score) | 1.520 (1.123 - 2.056) | **6.72E-03** |
| *MUC4* mutation (Yes vs. No) | 0.554 (0.359 - 0.854) | **7.59E-03** |

| **Renal papillary cell** | **Univariate** |  |
| --- | --- | --- |
| Signature 1 (high vs. low score) | 0.370 (0.157 - 0.871) | **0.023** |
| Signature 2 (high vs. low score) | 3.862 (1.565 - 9.526) | **3.36E-03** |
| TNM staging (II & III vs. I) | 2.71 (1.893 - 3.878) | **5.03E-08** |
|  | **Multivariate (Signature 1)** |  |
| Signature 1 (high vs. low score) | 0.461 (0.194 - 0.981) | **0.048** |
| TNM staging (II & III vs. I) | 2.621 (1.821 - 3.772) | **2.13E-07** |
|  | **Multivariate (Signature 2)** |  |
| Signature 2 (high vs. low score) | 3.057 (1.232 - 7.584) | **0.016** |
| TNM staging (II & III vs. I) | 2.616 (1.805 - 3.790) | **3.78E-07** |

| **Liver #1 (LIHC)** | **Univariate** |  |
| --- | --- | --- |
| Signature 1 (high vs. low score) | 0.656 (0.424 - 0.915) | **0.048** |
| TNM staging (II & III vs. I) | 2.000 (1.533 - 2.608) | **3.14E-07** |
|  | **Multivariate (Signature 1)** |  |
| Signature 1 (high vs. low score) | 0.717 (0.464 - 1.108) | 0.13 |
| TNM staging (II & III vs. I) | 1.999 (1.570 - 2.546) | **1.99E-08** |

| **Liver #2 (LIRI-JP)** | **Univariate** |  |
| --- | --- | --- |
| Signature 1 (high vs. low score) | 0.490 (0.259 - 0.938) | **0.031** |
| Signature 2 (high vs. low score) | 5.271 (2.429 - 11.44) | **2.61E-05** |
| Tumour size (> 3cm vs. < 3cm) | 2.415 (1.263 - 4.615) | **7.60E-03** |
| TNM staging (II & III vs. I) | 2.267 (1.556 - 3.304) | **2.03E-05** |
|  | **Multivariate (Signature 1)** |  |
| Signature 1 (high vs. low score) | 0.541 (0.283 - 0.904) | **0.043** |
| Tumour size (> 3cm vs. < 3cm) | 1.003 (0.997 - 1.009) | 0.38 |
| TNM staging (II & III vs. I) | 2.065 (1.374 - 3.104) | **0.000483** |
|  | **Multivariate (Signature 2)** |  |
| Signature 2 (high vs. low score) | 4.539 (2.055 - 10.029) | **1.84E-04** |
| Tumour size (> 3cm vs. < 3cm) | 1.001 (0.994 - 1.008) | 0.78 |
| TNM staging (II & III vs. I) | 1.926 (1.296 - 2.862) | **1.19E-03** |

| **Liver #3 (GSE14520)** | **Univariate** |  |
| --- | --- | --- |
| Signature 2 (high vs. low score) | 2.285 (1.458 - 3.580) | **3.12E-04** |
| Tumour size (> 5cm vs. < 5cm) | 2.159 (1.406 - 3.316) | **4.40E-04** |
| Cirrhosis (Yes vs. No) | 4.665 (1.147 - 18.97) | **0.031** |
| TNM staging (II & III vs. I) | 2.26 (1.706 - 2.994) | **1.30E-08** |
| BCLC staging (B & C vs. A & 0) | 2.181 (1.722 - 2.762) | **9.86E-11** |
| AFP (> 300ng/mL vs. < 300ng/mL) | 1.606 (1.049 - 2.46) | **0.029** |
|  | **Multivariate (Signature 2)** |  |
| Signature 2 (high vs. low score) | 2.012 (1.267 - 3.195) | **3.04E-03** |
| Tumour size (> 5cm vs. < 5cm) | 0.983 (0.589 - 1.642) | 0.94 |
| Cirrhosis (Yes vs. No) | 3.187 (0.762 - 13.335) | 0.11 |
| TNM staging (II & III vs. I) | 1.346 (0.914 - 1.980) | 0.13 |
| BCLC staging (B & C vs. A & 0) | 1.873 (1.332 - 2.634) | **3.08E-04** |

| **Lung** | **Univariate** |
| --- | --- |
| Signature 1 (high vs. low score) | 0.6245 (0.443 - 0.879) | **6.97E-03** |
| Signature 2 (high vs. low score) | 1.562 (1.116 - 2.188) | **9.44E-03** |
| TNM staging (II & III vs. I) | 1.597 (1.364 - 1.870) | **6.11E-09** |
| **Multivariate (Signature 1)** |  |  |
| Signature 1 (high vs. low score) | 0.715 (0.505 - 0.913) | **0.039** |
| TNM staging (II & III vs. I) | 1.553 (1.324 - 1.822) | **6.74E-08** |
| **Multivariate (Signature 2)** |  |  |
| Signature 2 (high vs. low score) | 1.496 (1.067 - 2.097) | **0.019** |
| TNM staging (II & III vs. I) | 1.587 (1.353 - 1.861) | **1.36E-08** |
| **Pancreas** |  |  |
| **Univariate** |  |  |
| Signature 1 (high vs. low score) | 0.454 (0.278 - 0.741) | **1.59E-03** |
| Signature 2 (high vs. low score) | 1.969 (1.217 - 3.186) | **5.81E-03** |
| TNM staging (II & III vs. I) | 1.339 (0.897 - 1.998) | 0.153 |
| **Multivariate (Signature 1)** |  |  |
| Signature 1 (high vs. low score) | 0.472 (0.284 - 0.783) | **3.62E-03** |
| TNM staging (II & III vs. I) | 1.150 (0.722 - 1.832) | 0.56 |
| **Multivariate (Signature 2)** |  |  |
| Signature 2 (high vs. low score) | 1.888 (1.157 - 3.082) | **0.011** |
| TNM staging (II & III vs. I) | 1.205 (0.785 - 1.851) | 0.394 |
| **Stomach** |  |  |
| **Univariate** |  |  |
| Signature 2 (high vs. low score) | 1.725 (1.142 - 2.605) | **9.50E-03** |
| TNM staging (II & III vs. I) | 1.372 (1.067 - 1.765) | **0.013** |
| *PCDHA1* mutation (Yes vs. No) | 1.525 (1.007 - 2.307) | **0.046** |
| *PCDHA2* mutation (Yes vs. No) | 1.604 (1.061 - 2.427) | **0.025** |
| *TP53* mutation (Yes vs. No) | 1.296 (0.853 - 1.967) | 0.22 |
| *TTN* mutation (Yes vs. No) | 1.115 (0.740 - 1.682) | 0.60 |
| **Multivariate (Signature 2)** |  |  |
| Signature 2 (high vs. low score) | 1.800 (1.184 - 2.737) | **5.90E-03** |
| TNM staging (II & III vs. I) | 1.464 (1.130 - 1.898) | **3.90E-03** |
| *PCDHA1* mutation (Yes vs. No) | 1.501 (0.988 - 2.281) | 0.056 |
| *PCDHA2* mutation (Yes vs. No) | 1.558 (1.026 - 2.365) | **0.037** |
| **Endometrial** |  |  |
| **Univariate** |  |  |
| Signature 1 (high vs. low score) | 0.401 (0.229 - 0.702) | **1.39E-03** |
| TNM staging (II & III vs. I) | 1.802 (1.431 - 2.269) | **5.69E-07** |
| *PCDHA1* mutation (Yes vs. No) | 0.516 (0.272 - 0.978) | **0.042** |
| *PIK3CA* mutation (Yes vs. No) | 0.362 (0.190 - 0.689) | **1.98E-03** |
| *PIK3R1* mutation (Yes vs. No) | 1.180 (0.645 - 2.158) | 0.592 |
| *PTEN* mutation (Yes vs. No) | 0.427 (0.234 - 0.781) | **5.75E-03** |
| *TP53* mutation (Yes vs. No) | 1.780 (1.025 - 3.090) | **0.041** |
| **Multivariate (Signature 1)** |  |  |
| Signature 1 (high vs. low score) | 0.519 (0.293 - 0.920) | **0.024** |
| TNM staging (II & III vs. I) | 1.589 (1.245 - 2.028) | **2.02E-04** |
| *PCDHA1* mutation (Yes vs. No) | 0.803 (0.376 - 1.712) | 0.56 |
| *PIK3CA* mutation (Yes vs. No) | 0.428 (0.216 - 0.848) | **0.015** |
| *PTEN* mutation (Yes vs. No) | 0.904 (0.427 - 1.910) | 0.79 |
| *TP53* mutation (Yes vs. No) | 1.567 (0.867 - 2.832) | 0.13 |

**Table S4.**
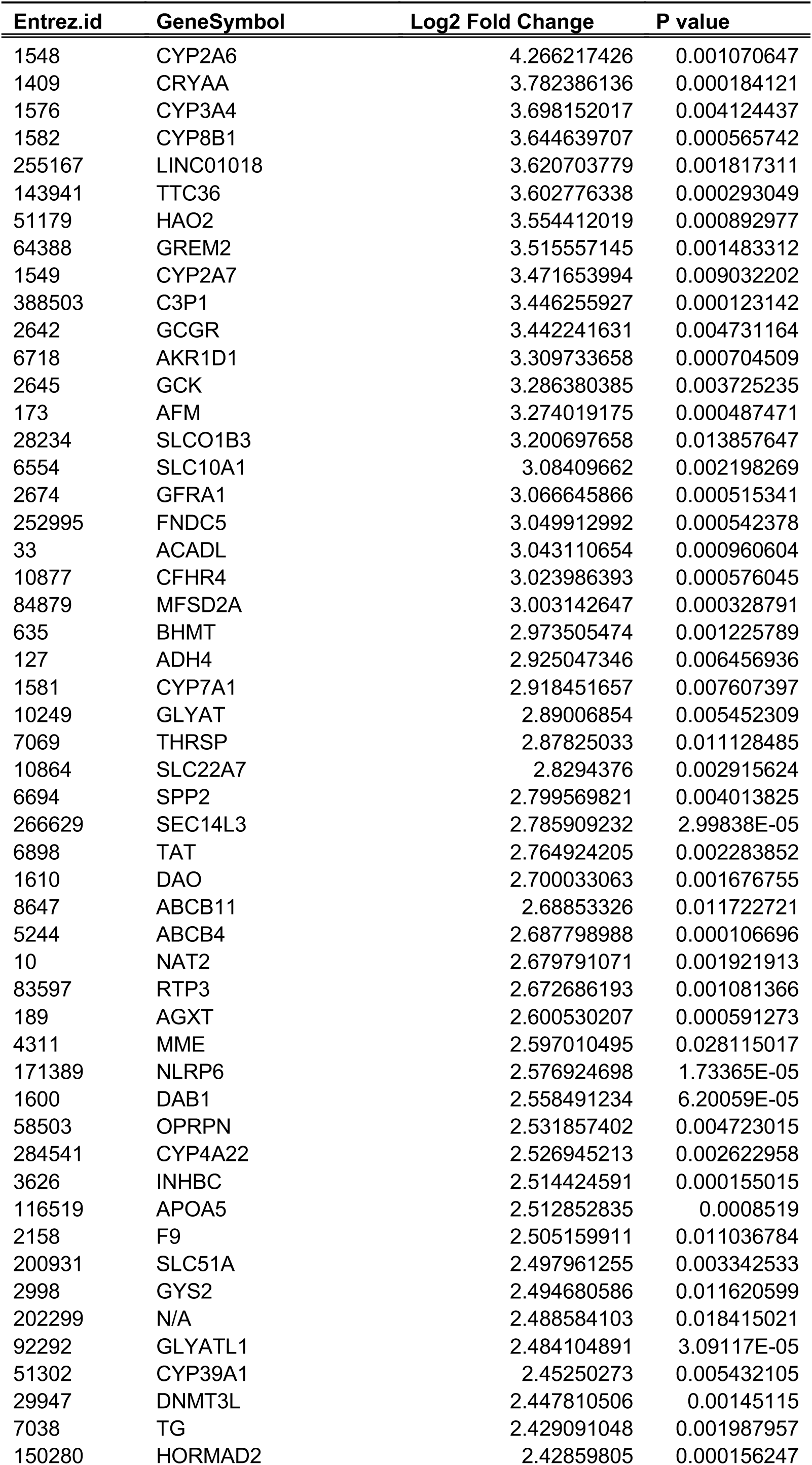
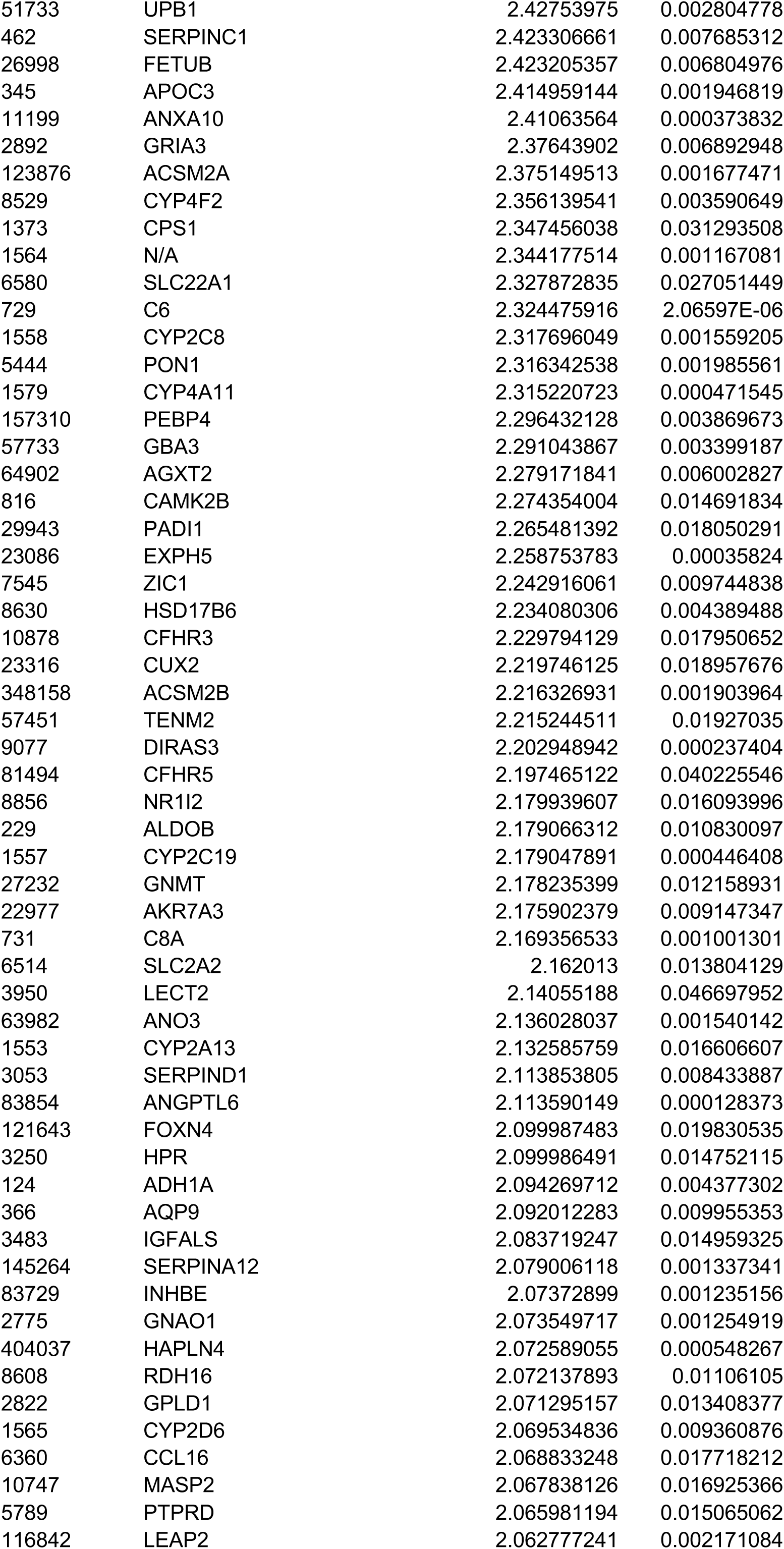
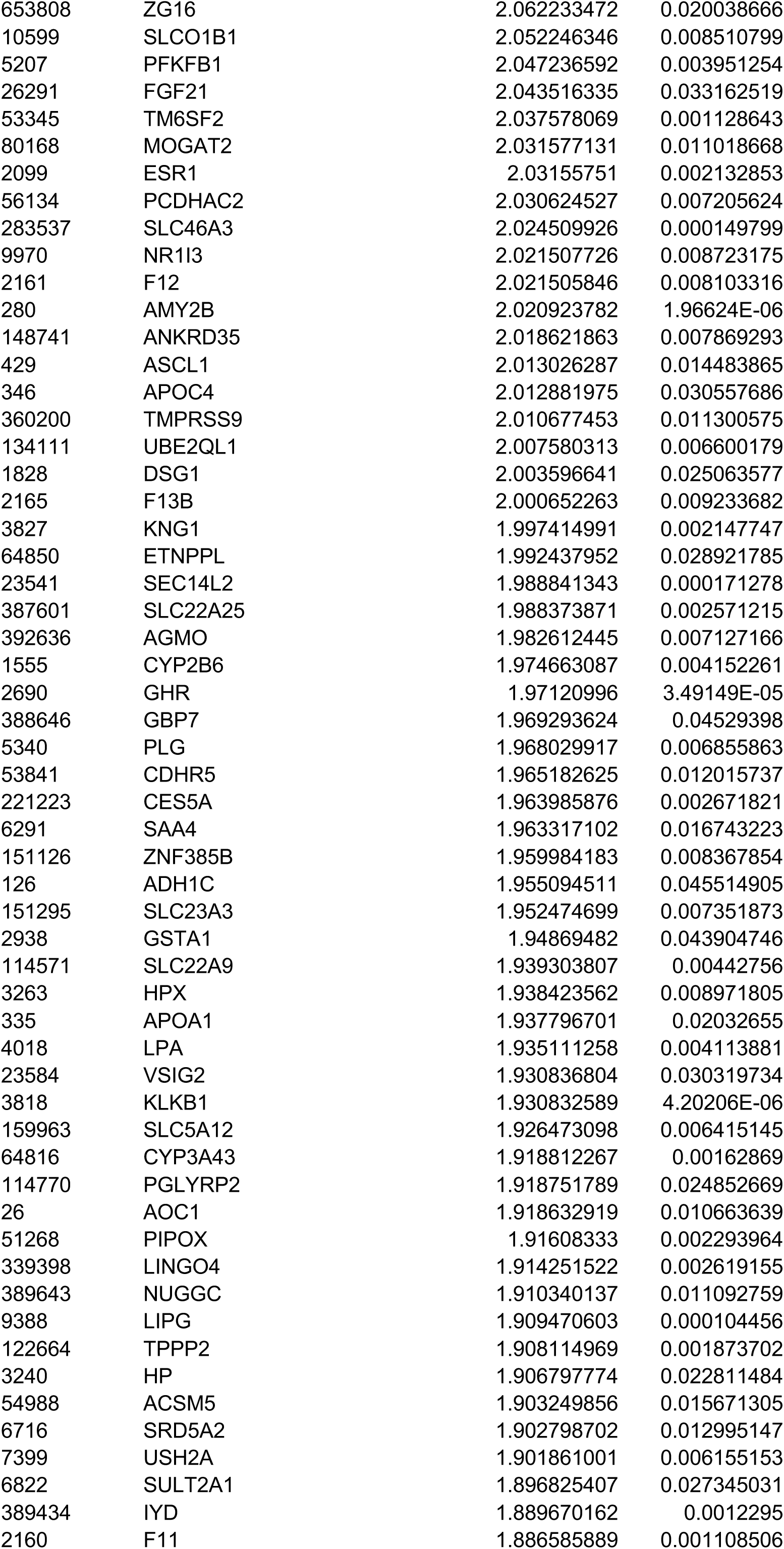
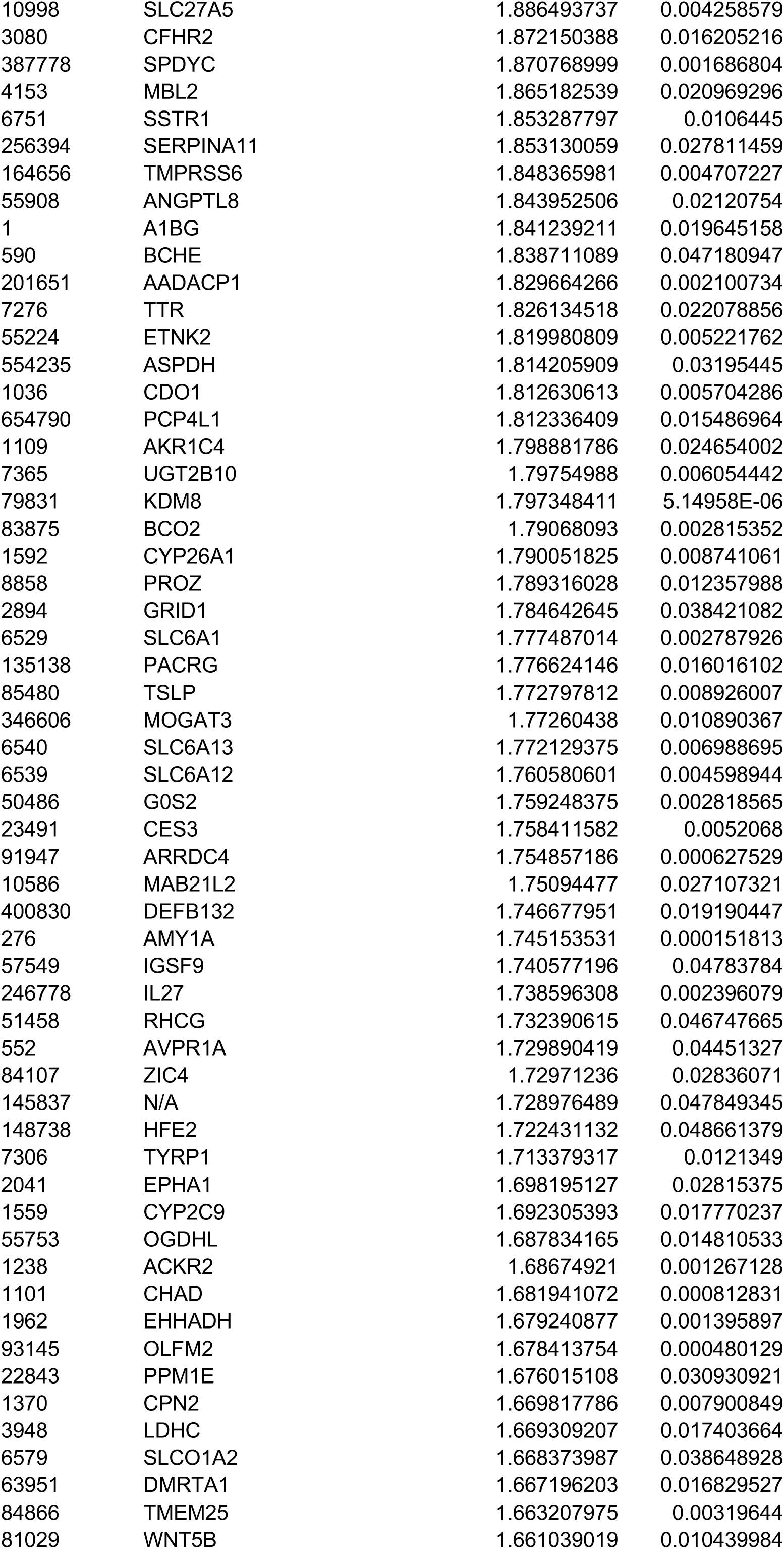
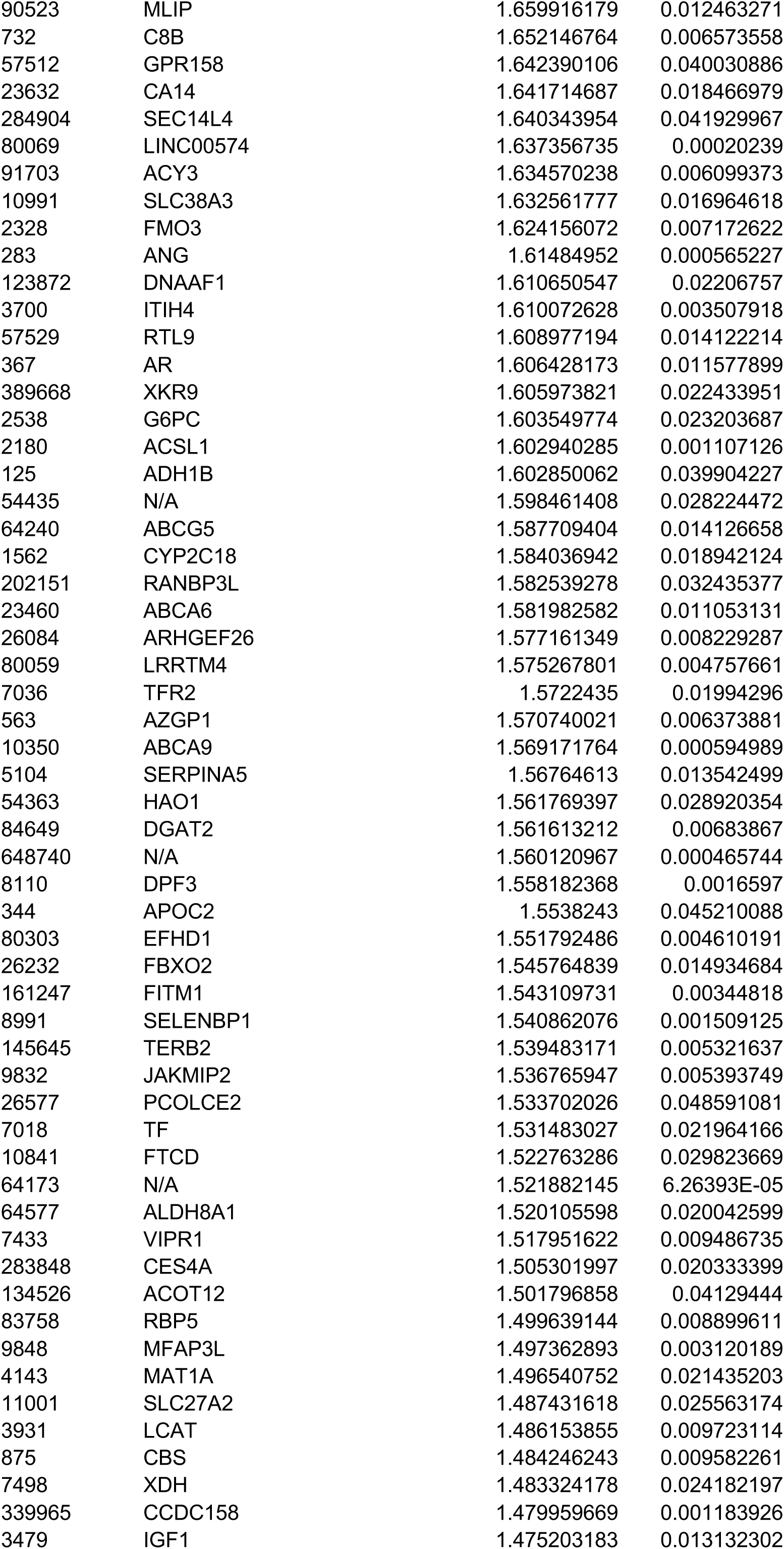
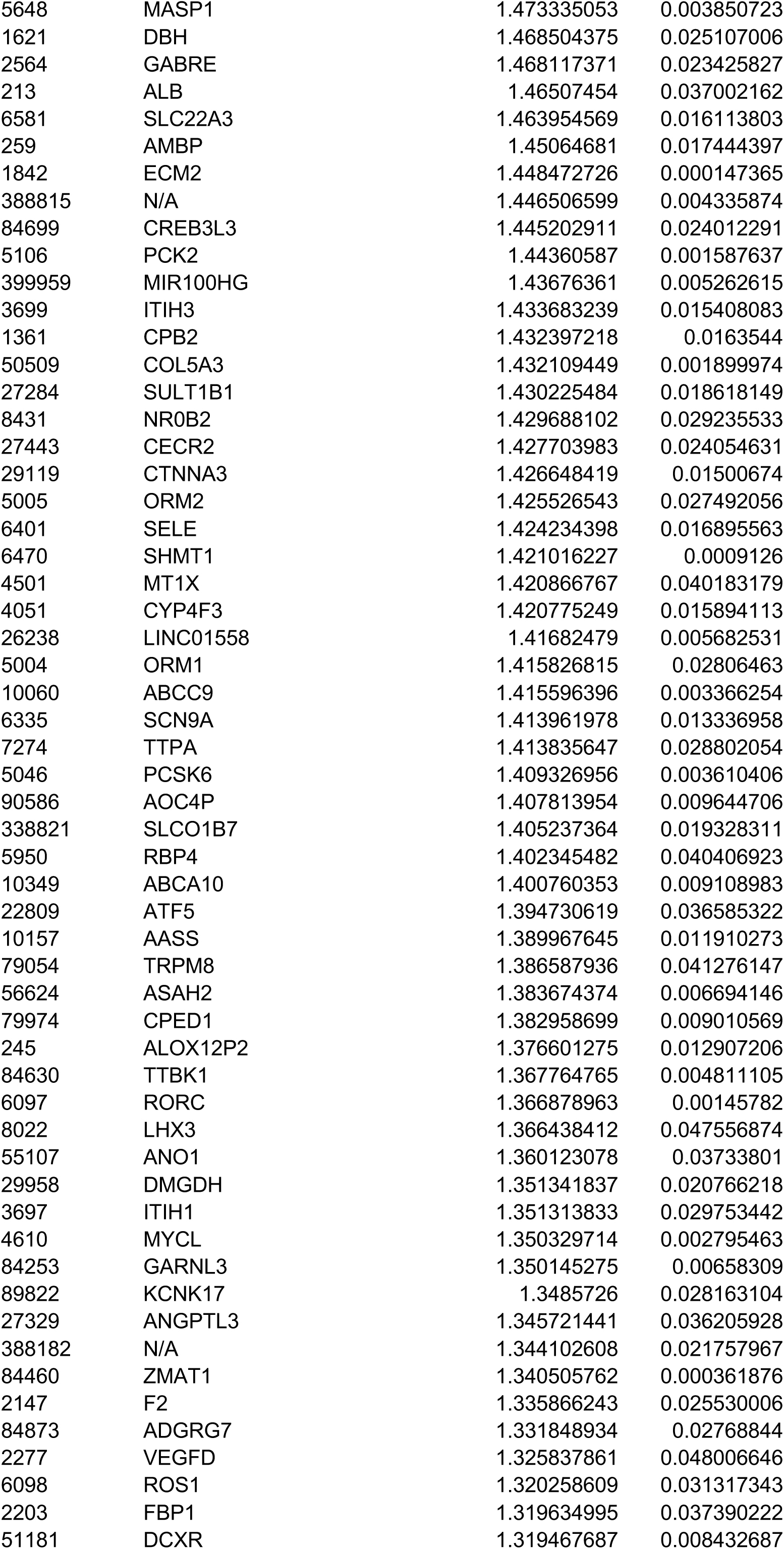
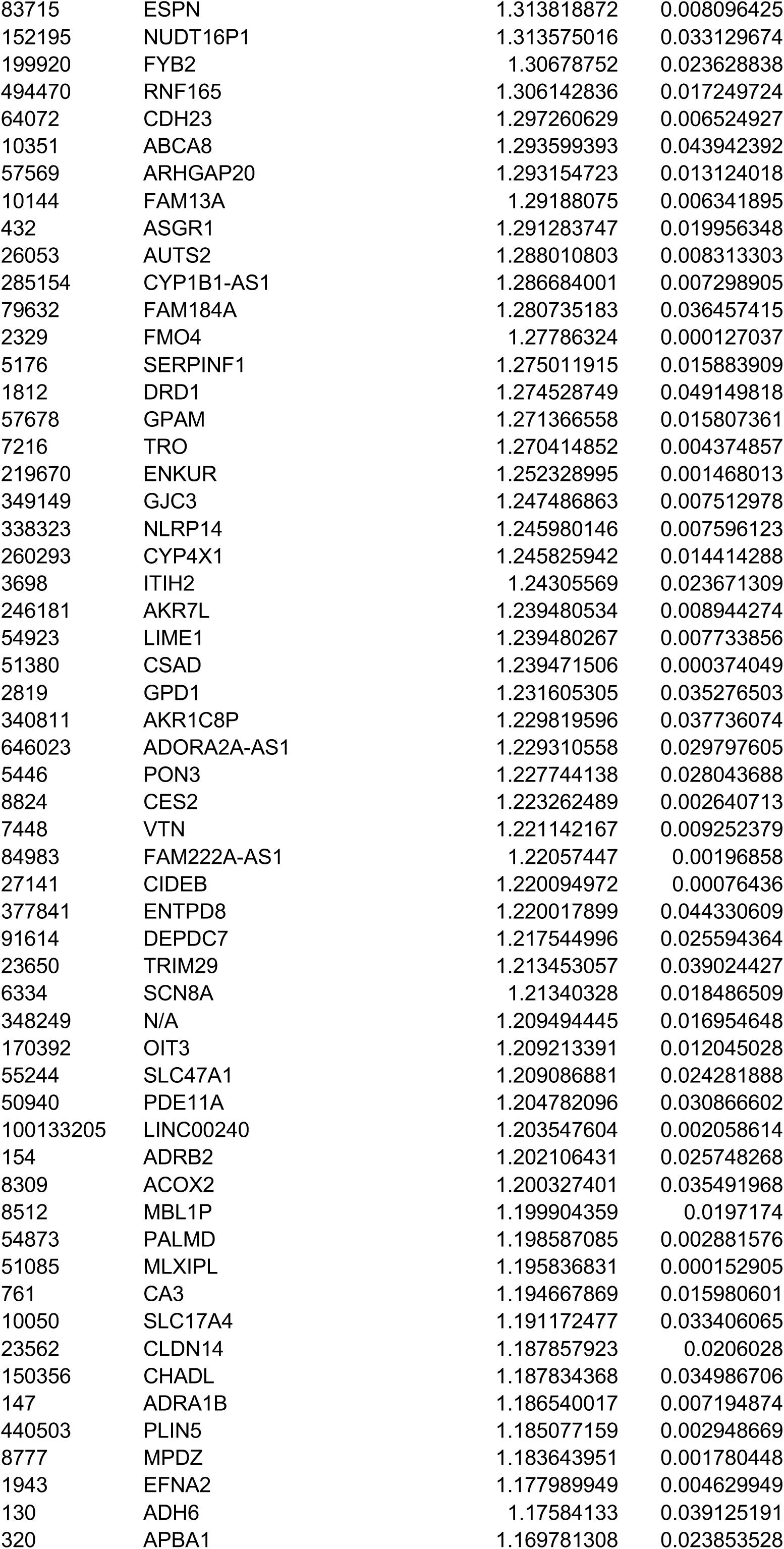
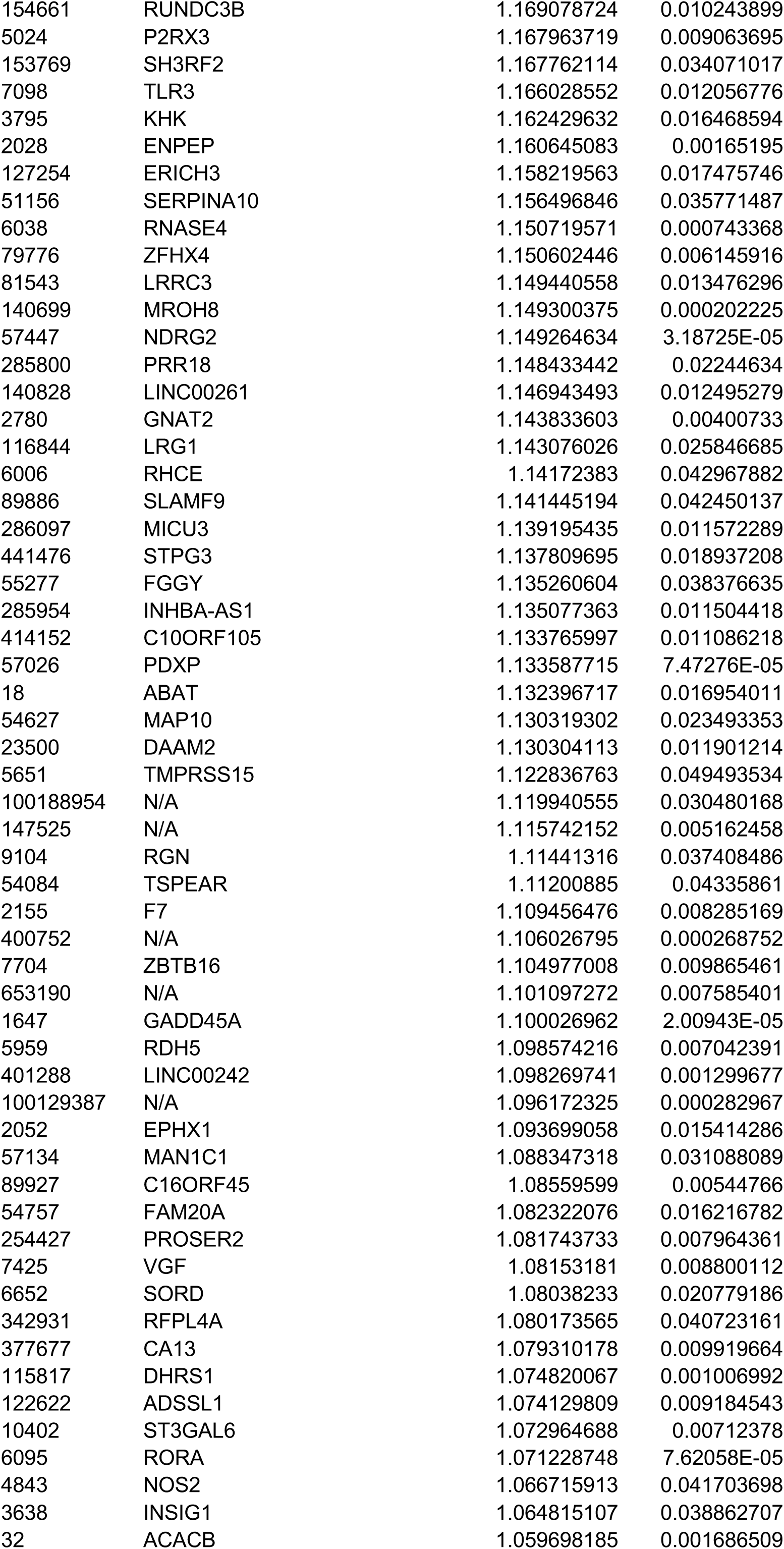
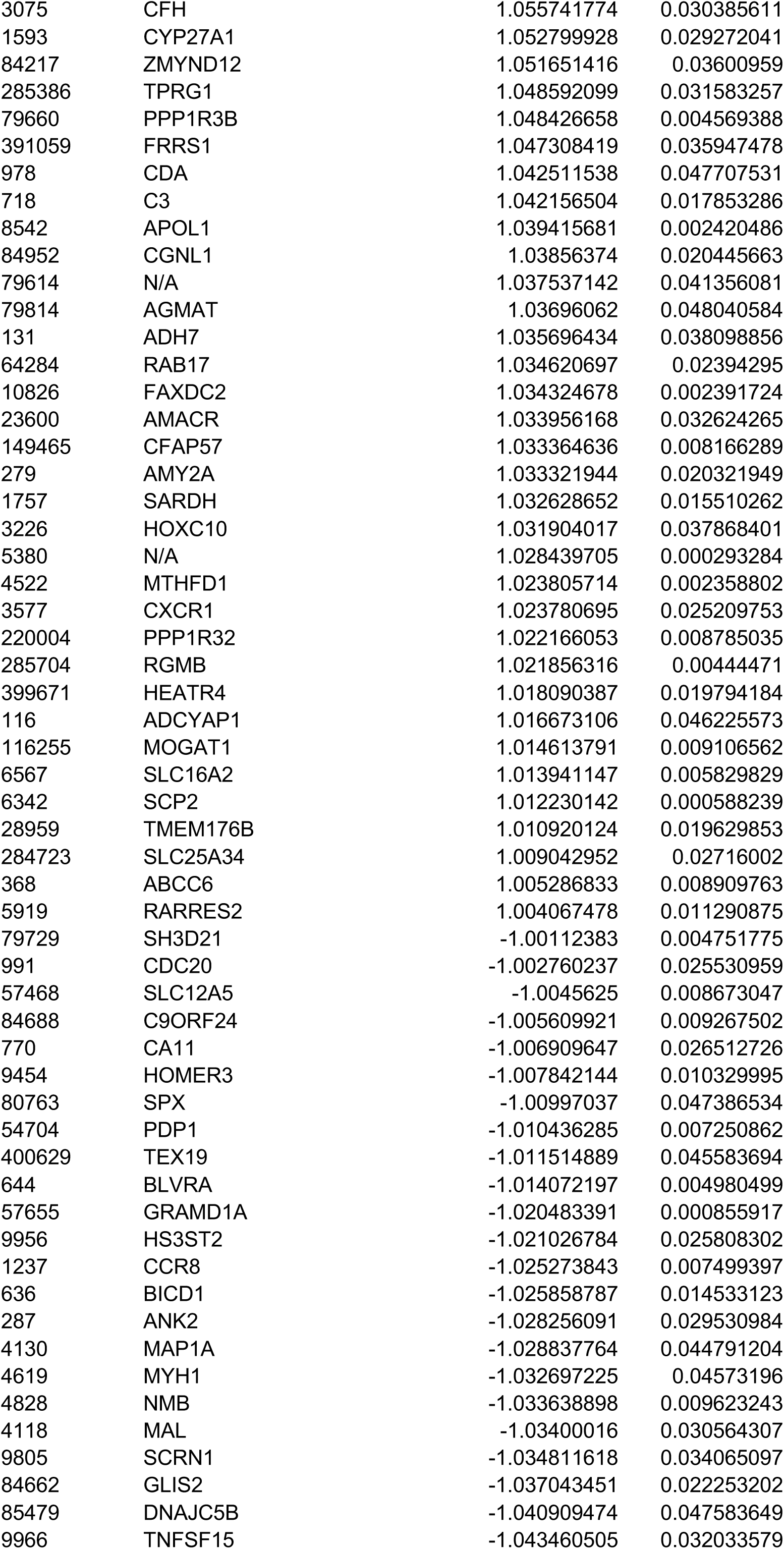
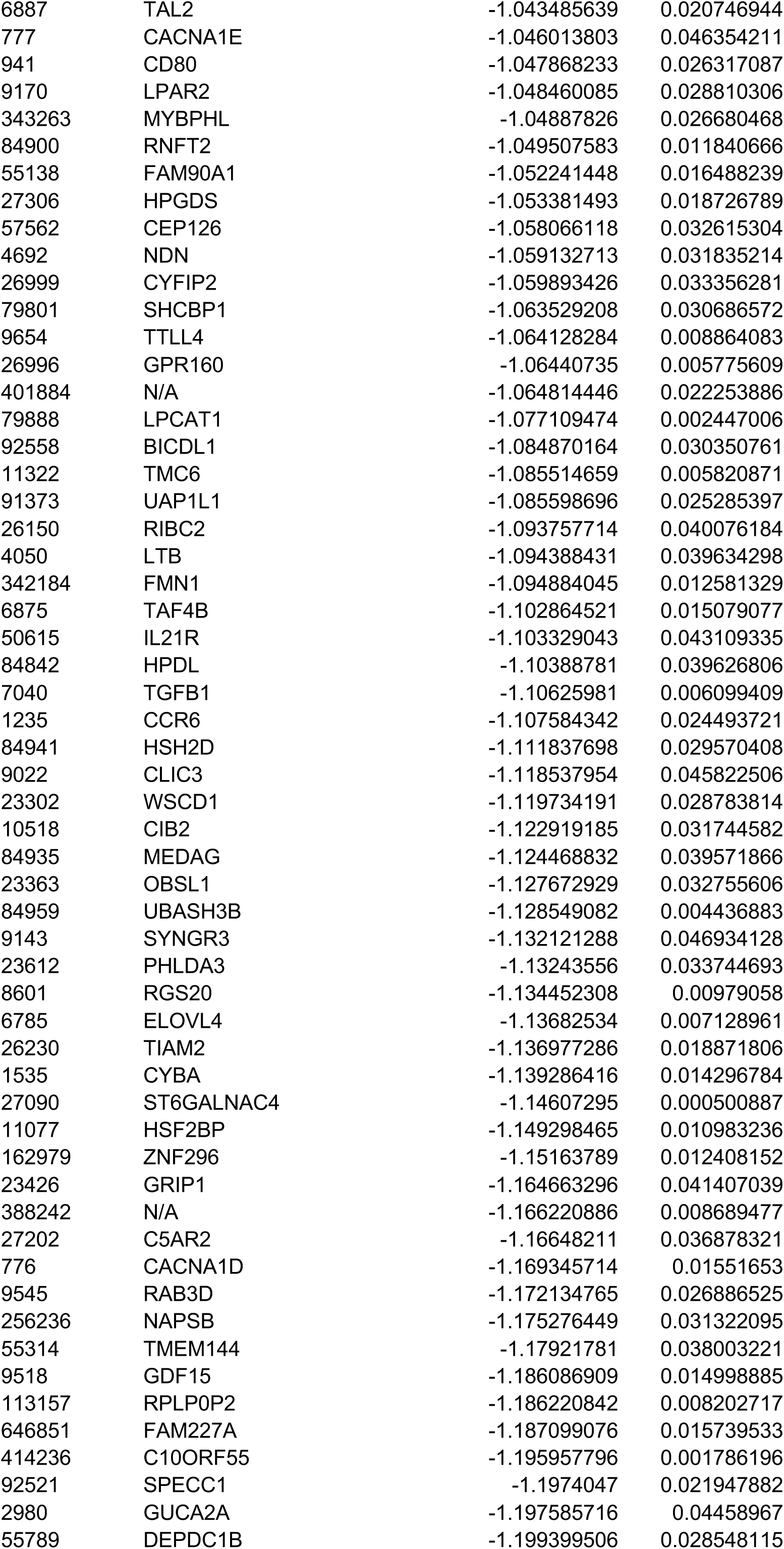
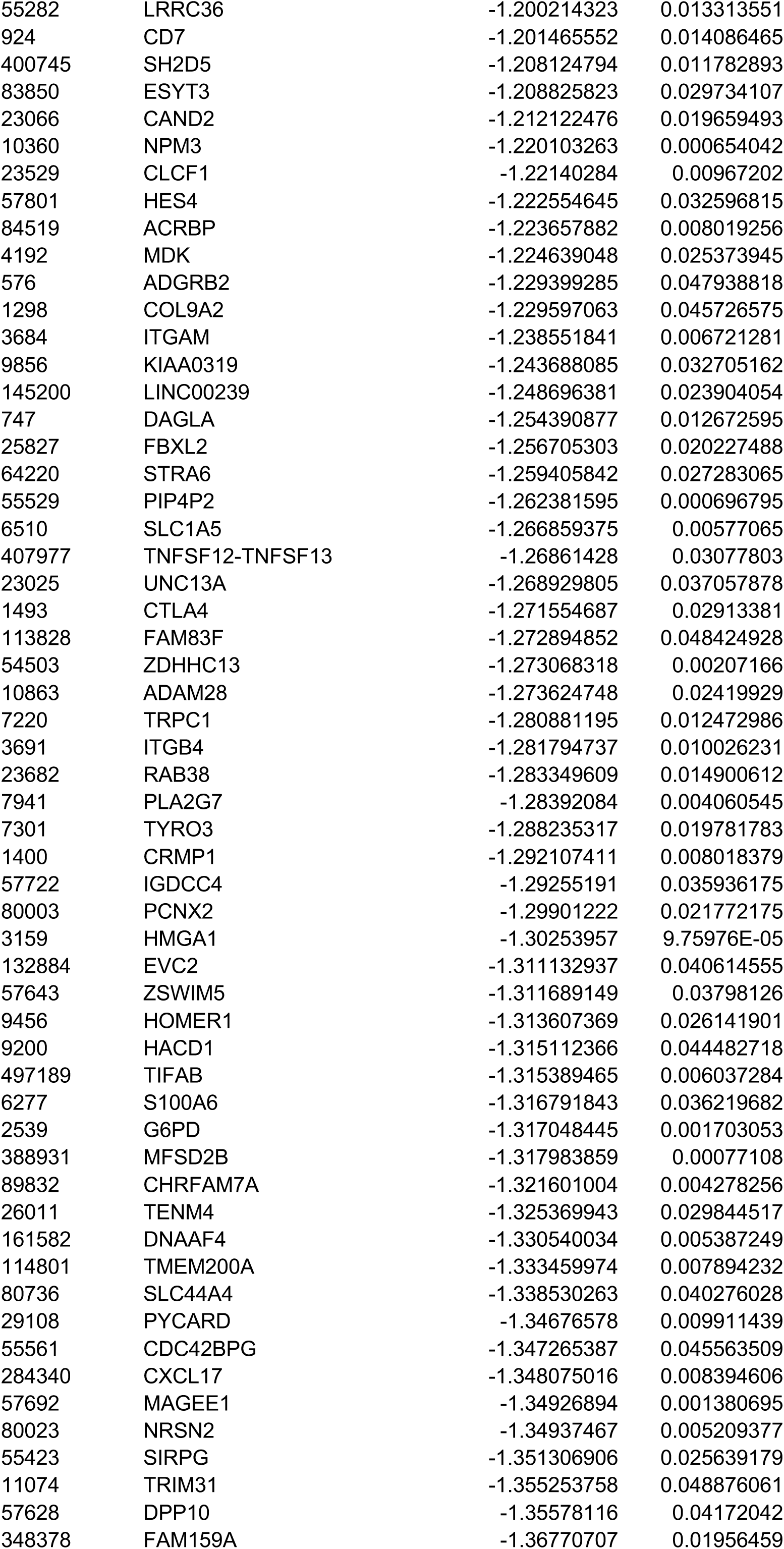
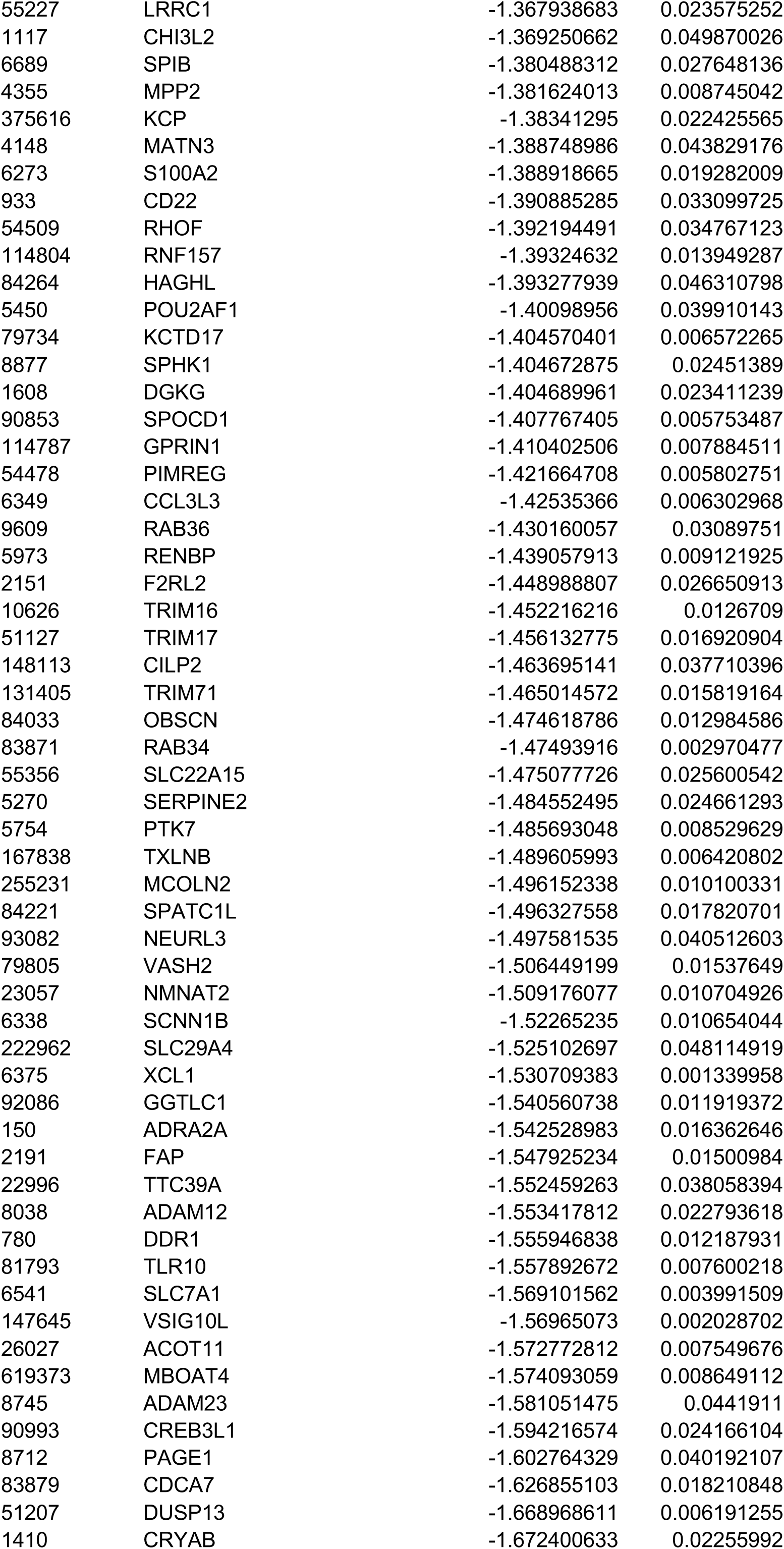
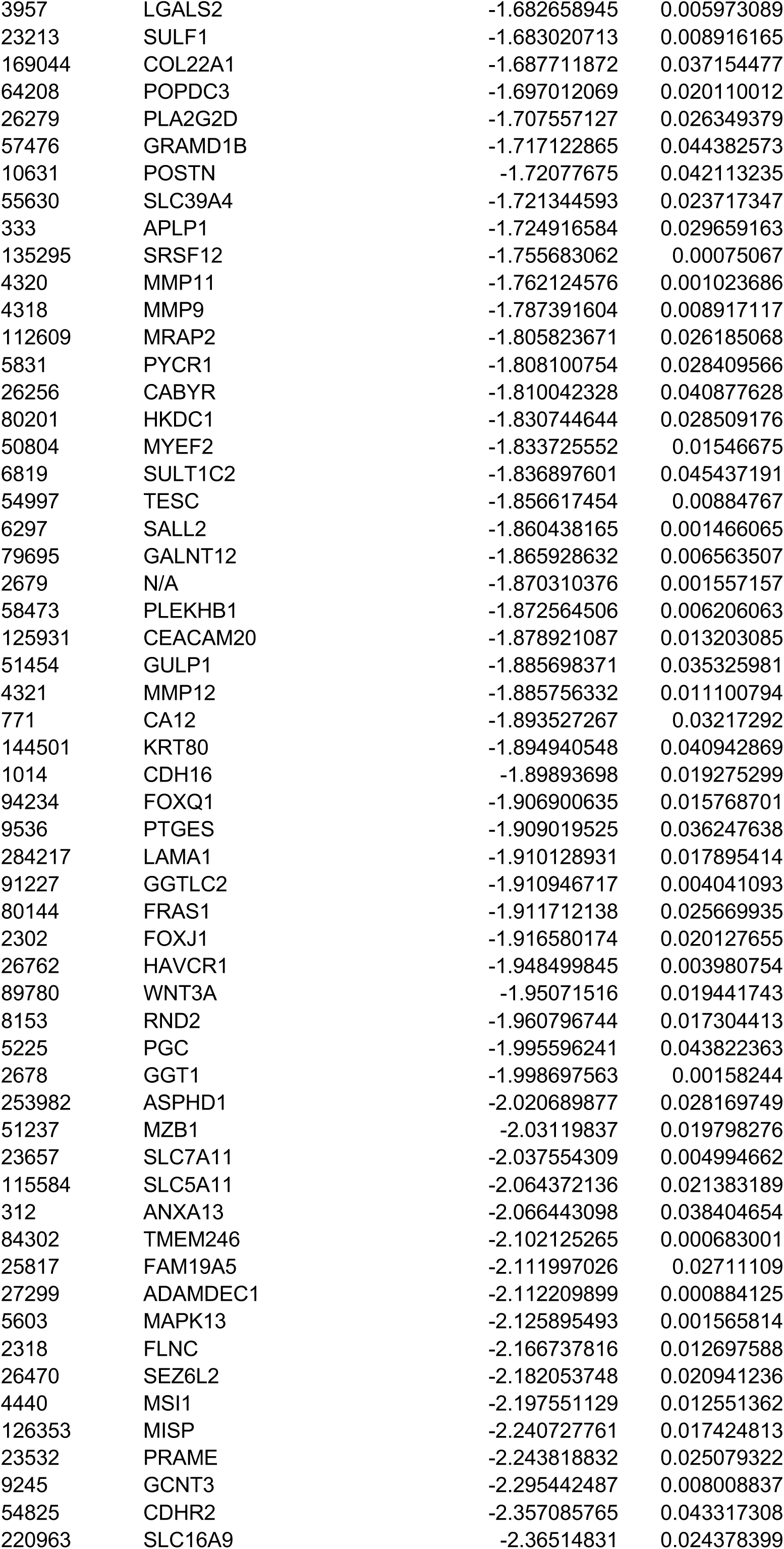
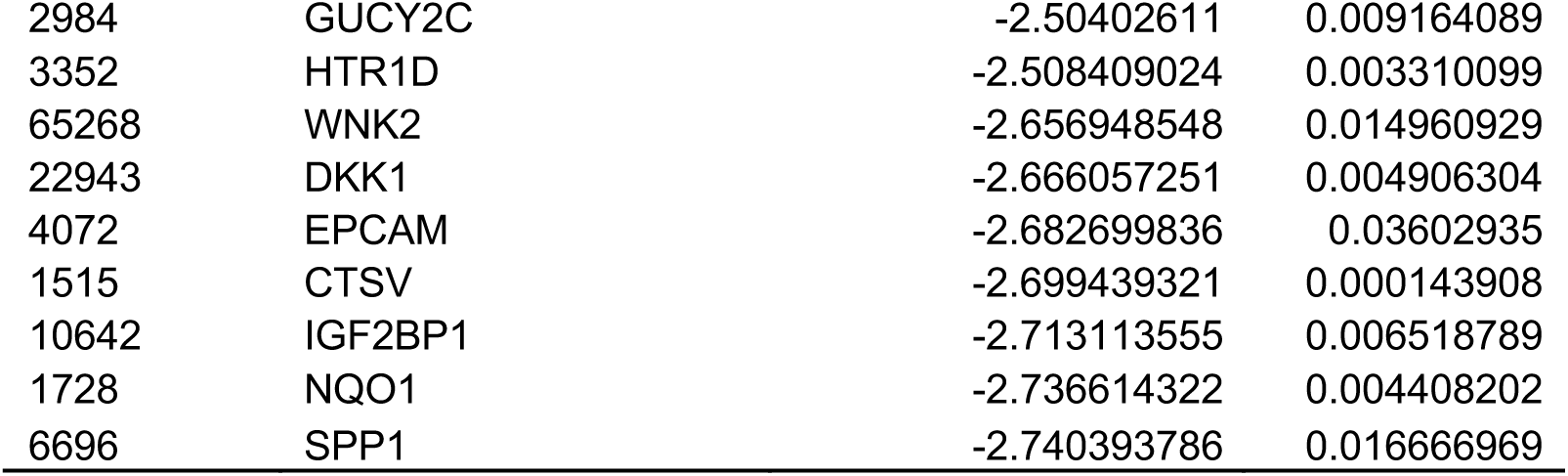
Differentially expressed genes between *KDM8*-high and -low groups in hepatocellular carcinoma (LIHC cohort).

**Table S5.** Significantly enriched biological pathways of differentially expressed genes.

| Category | Ontology | P Value | Fold Enrichment |
| --- | --- | --- | --- |
| GO_biological_processes | GO:0006805~xenobiotic metabolic process | 3.46E-17 | 8.87 |
| GO_biological_processes | GO:0055114~oxidation-reduction process | 5.66E-15 | 2.97 |
| KEGG_pathway | hsa01100:Metabolic pathways | 1.26E-13 | 1.96 |
| KEGG_pathway | hsa04610:Complement and coagulation cascades | 1.18E-11 | 6.68 |
| GO_biological_processes | GO:0006699~bile acid biosynthetic process | 4.77E-11 | 15.21 |
| KEGG_pathway | hsa00830:Retinol metabolism | 2.51E-09 | 6.07 |
| GO_biological_processes | GO:0010951~negative regulation of endopeptidase activity | 4.15E-09 | 4.84 |
| GO_biological_processes | GO:0019373~epoxygenase P450 pathway | 5.00E-09 | 14.78 |
| KEGG_pathway | hsa05204:Chemical carcinogenesis | 1.15E-08 | 5.21 |
| KEGG_pathway | hsa00120:Primary bile acid biosynthesis | 1.30E-08 | 12.90 |
| KEGG_pathway | hsa00980:Metabolism of xenobiotics by cytochrome P450 | 2.12E-08 | 5.34 |
| GO_biological_processes | GO:0008202~steroid metabolic process | 3.70E-08 | 8.05 |
| KEGG_pathway | hsa00982:Drug metabolism - cytochrome P450 | 3.87E-08 | 5.48 |
| KEGG_pathway | hsa04976:Bile secretion | 4.83E-08 | 5.40 |
| GO_biological_processes | GO:0042738~exogenous drug catabolic process | 6.80E-08 | 17.74 |
| GO_biological_processes | GO:0006869~lipid transport | 1.18E-07 | 5.60 |
| GO_biological_processes | GO:0070328~triglyceride homeostasis | 2.45E-07 | 10.24 |
| GO_biological_processes | GO:0017144~drug metabolic process | 3.56E-07 | 9.86 |
| GO_biological_processes | GO:0055085~transmembrane transport | 5.19E-07 | 3.05 |
| GO_biological_processes | GO:0015721~bile acid and bile salt transport | 4.43E-06 | 8.87 |
| KEGG_pathway | hsa03320:PPAR signaling pathway | 7.57E-06 | 4.58 |
| GO_biological_processes | GO:0001523~retinoid metabolic process | 1.44E-05 | 5.24 |
| GO_biological_processes | GO:0006631~fatty acid metabolic process | 1.96E-05 | 5.63 |
| KEGG_pathway | hsa00071:Fatty acid degradation | 2.03E-05 | 5.48 |
| GO_biological_processes | GO:0002576~platelet degranulation | 2.99E-05 | 3.88 |
| GO_biological_processes | GO:0019432~triglyceride biosynthetic process | 3.57E-05 | 8.19 |
| KEGG_pathway | hsa00010:Glycolysis / Gluconeogenesis | 3.96E-05 | 4.26 |
| GO_biological_processes | GO:0006069~ethanol oxidation | 4.65E-05 | 13.31 |
| KEGG_pathway | hsa00260:Glycine, serine and threonine metabolism | 4.67E-05 | 5.62 |
| GO_biological_processes | GO:0042632~cholesterol homeostasis | 1.25E-04 | 4.57 |
| KEGG_pathway | hsa00350:Tyrosine metabolism | 1.33E-04 | 5.64 |
| GO_biological_processes | GO:0051919~positive regulation of fibrinolysis | 2.04E-04 | 26.61 |
| GO_biological_processes | GO:0097267~omega-hydroxylase P450 pathway | 2.13E-04 | 14.78 |
| GO_biological_processes | GO:0009636~response to toxic substance | 3.26E-04 | 3.76 |
| KEGG_pathway | hsa04146:Peroxisome | 3.33E-04 | 3.44 |
| GO_biological_processes | GO:0045723~positive regulation of fatty acid biosynthetic process | 3.44E-04 | 13.31 |
| GO_biological_processes | GO:0043691~reverse cholesterol transport | 4.17E-04 | 8.87 |
| GO_biological_processes | GO:0007597~blood coagulation, intrinsic pathway | 4.17E-04 | 8.87 |
| KEGG_pathway | hsa00590:Arachidonic acid metabolism | 4.26E-04 | 3.89 |
| GO_biological_processes | GO:0042157~lipoprotein metabolic process | 4.63E-04 | 5.60 |
| GO_biological_processes | GO:0051180~vitamin transport | 4.97E-04 | 21.29 |
| GO_biological_processes | GO:0006071~glycerol metabolic process | 5.25E-04 | 12.10 |
| GO_biological_processes | GO:0006953~acute-phase response | 5.46E-04 | 5.46 |
| KEGG_pathway | hsa00500:Starch and sucrose metabolism | 5.75E-04 | 5.32 |
| KEGG_pathway | hsa01130:Biosynthesis of antibiotics | 5.96E-04 | 2.28 |
| GO_biological_processes | GO:0006810~transport | 6.94E-04 | 2.06 |
| KEGG_pathway | hsa02010:ABC transporters | 7.03E-04 | 4.49 |
| GO_biological_processes | GO:0030449~regulation of complement activation | 7.48E-04 | 6.21 |
| GO_biological_processes | GO:0042730~fibrinolysis | 9.03E-04 | 7.60 |
| GO_biological_processes | GO:0016098~monoterpenoid metabolic process | 9.66E-04 | 17.74 |
| GO_biological_processes | GO:0042737~drug catabolic process | 9.66E-04 | 17.74 |
| GO_biological_processes | GO:0006957~complement activation, alternative pathway | 0.001070284 | 10.24 |
| GO_biological_processes | GO:0006637~acyl-CoA metabolic process | 0.001132542 | 7.26 |
| GO_biological_processes | GO:0043252~sodium-independent organic anion transport | 0.001402936 | 6.94 |
| GO_biological_processes | GO:0051289~protein homotetramerization | 0.001717738 | 3.99 |
| GO_biological_processes | GO:0015695~organic cation transport | 0.001924609 | 8.87 |
| GO_biological_processes | GO:0008209~androgen metabolic process | 0.00249067 | 8.32 |
| GO_biological_processes | GO:0010898~positive regulation of triglyceride catabolic process | 0.00255538 | 13.31 |
| GO_biological_processes | GO:0006633~fatty acid biosynthetic process | 0.003122356 | 4.09 |
| GO_biological_processes | GO:0055091~phospholipid homeostasis | 0.003726728 | 11.83 |
| GO_biological_processes | GO:0031638~zymogen activation | 0.003726728 | 11.83 |
| GO_biological_processes | GO:0007596~blood coagulation | 0.004023522 | 2.31 |
| GO_biological_processes | GO:0050746~regulation of lipoprotein metabolic process | 0.004111183 | 26.61 |
| GO_biological_processes | GO:0030573~bile acid catabolic process | 0.004111183 | 26.61 |
| KEGG_pathway | hsa00983:Drug metabolism - other enzymes | 0.004345618 | 3.82 |
| GO_biological_processes | GO:0008152~metabolic process | 0.004412615 | 2.38 |
| GO_biological_processes | GO:0042593~glucose homeostasis | 0.004607137 | 2.90 |
| GO_biological_processes | GO:0042572~retinol metabolic process | 0.004786067 | 5.32 |
| GO_biological_processes | GO:0006730~one-carbon metabolic process | 0.004786067 | 5.32 |
| GO_biological_processes | GO:0006855~drug transmembrane transport | 0.004851097 | 7.00 |
| KEGG_pathway | hsa01200:Carbon metabolism | 0.005030137 | 2.52 |
| KEGG_pathway | hsa04922:Glucagon signaling pathway | 0.005057234 | 2.66 |
| GO_biological_processes | GO:0051156~glucose 6-phosphate metabolic process | 0.005176482 | 10.64 |
| GO_biological_processes | GO:0046470~phosphatidylcholine metabolic process | 0.005176482 | 10.64 |
| GO_biological_processes | GO:0006000~fructose metabolic process | 0.005176482 | 10.64 |
| GO_biological_processes | GO:0031667~response to nutrient levels | 0.005533575 | 5.15 |
| GO_biological_processes | GO:0006094~gluconeogenesis | 0.005710899 | 4.23 |
| KEGG_pathway | hsa00910:Nitrogen metabolism | 0.006220473 | 6.45 |
| GO_biological_processes | GO:0007584~response to nutrient | 0.006410732 | 3.24 |
| GO_biological_processes | GO:0017187~peptidyl-glutamic acid carboxylation | 0.00692097 | 9.68 |
| GO_biological_processes | GO:0010867~positive regulation of triglyceride biosynthetic process | 0.00692097 | 9.68 |
| GO_biological_processes | GO:0032094~response to food | 0.007059136 | 6.34 |
| GO_biological_processes | GO:0001676~long-chain fatty acid metabolic process | 0.007059136 | 6.34 |
| GO_biological_processes | GO:0006651~diacylglycerol biosynthetic process | 0.008017984 | 19.96 |
| GO_biological_processes | GO:0019448~L-cysteine catabolic process | 0.008017984 | 19.96 |
| GO_biological_processes | GO:0036101~leukotriene B4 catabolic process | 0.008017984 | 19.96 |
| GO_biological_processes | GO:0010897~negative regulation of triglyceride catabolic process | 0.008017984 | 19.96 |
| GO_biological_processes | GO:0015711~organic anion transport | 0.008973455 | 8.87 |
| GO_biological_processes | GO:0030212~hyaluronan metabolic process | 0.008973455 | 8.87 |
| GO_biological_processes | GO:0070989~oxidative demethylation | 0.008973455 | 8.87 |
| GO_biological_processes | GO:0007586~digestion | 0.009074707 | 3.38 |
| GO_biological_processes | GO:0042594~response to starvation | 0.009351338 | 4.56 |
| KEGG_pathway | hsa05150:Staphylococcus aureus infection | 0.010557477 | 3.25 |
| GO_biological_processes | GO:0051384~response to glucocorticoid | 0.010712715 | 3.28 |
| GO_biological_processes | GO:0042493~response to drug | 0.01097762 | 1.84 |
| KEGG_pathway | hsa04973:Carbohydrate digestion and absorption | 0.011053965 | 3.66 |
| GO_biological_processes | GO:0019915~lipid storage | 0.011469433 | 5.54 |
| GO_biological_processes | GO:0007188~adenylate cyclase-modulating G-protein coupled receptor signaling pathway | 0.011811436 | 4.32 |
| GO_biological_processes | GO:0007204~positive regulation of cytosolic calcium ion concentration | 0.01214725 | 2.38 |
| GO_biological_processes | GO:0015812~gamma-aminobutyric acid transport | 0.013032193 | 15.97 |
| GO_biological_processes | GO:0019439~aromatic compound catabolic process | 0.013032193 | 15.97 |
| GO_biological_processes | GO:0002682~regulation of immune system process | 0.013032193 | 15.97 |
| GO_biological_processes | GO:0033344~cholesterol efflux | 0.013257496 | 5.32 |
| KEGG_pathway | hsa00051:Fructose and mannose metabolism | 0.013760322 | 4.11 |
| GO_biological_processes | GO:0006508~proteolysis | 0.013913326 | 1.60 |
| GO_biological_processes | GO:0033700~phospholipid efflux | 0.014041733 | 7.60 |
| GO_biological_processes | GO:0042573~retinoic acid metabolic process | 0.014041733 | 7.60 |
| GO_biological_processes | GO:0007155~cell adhesion | 0.014451138 | 1.62 |
| GO_biological_processes | GO:0046487~glyoxylate metabolic process | 0.015213524 | 5.12 |
| GO_biological_processes | GO:0006629~lipid metabolic process | 0.015330745 | 2.20 |
| GO_biological_processes | GO:0006691~leukotriene metabolic process | 0.017071014 | 7.10 |
| GO_biological_processes | GO:0034375~high-density lipoprotein particle remodeling | 0.017071014 | 7.10 |
| GO_biological_processes | GO:0010575~positive regulation of vascular endothelial growth factor production | 0.017342128 | 4.93 |
| GO_biological_processes | GO:0006749~glutathione metabolic process | 0.018035879 | 3.33 |
| KEGG_pathway | hsa00232:Caffeine metabolism | 0.01878889 | 13.16 |
| GO_biological_processes | GO:0042426~choline catabolic process | 0.019065477 | 13.31 |
| GO_biological_processes | GO:0009804~coumarin metabolic process | 0.019065477 | 13.31 |
| GO_biological_processes | GO:0006739~NADP metabolic process | 0.019065477 | 13.31 |
| GO_biological_processes | GO:0046483~heterocycle metabolic process | 0.019065477 | 13.31 |
| GO_biological_processes | GO:0008206~bile acid metabolic process | 0.020435709 | 6.65 |
| GO_biological_processes | GO:0006898~receptor-mediated endocytosis | 0.023314405 | 2.00 |
| KEGG_pathway | hsa04931:Insulin resistance | 0.024468004 | 2.23 |
| GO_biological_processes | GO:0097264~self proteolysis | 0.026034557 | 11.41 |
| GO_biological_processes | GO:0010890~positive regulation of sequestering of triglyceride | 0.026034557 | 11.41 |
| GO_biological_processes | GO:0042908~xenobiotic transport | 0.026034557 | 11.41 |
| GO_biological_processes | GO:0015872~dopamine transport | 0.026034557 | 11.41 |
| GO_biological_processes | GO:0001867~complement activation, lectin pathway | 0.026034557 | 11.41 |
| GO_biological_processes | GO:0044539~long-chain fatty acid import | 0.026034557 | 11.41 |
| KEGG_pathway | hsa00270:Cysteine and methionine metabolism | 0.02751594 | 3.46 |
| GO_biological_processes | GO:0060384~innervation | 0.028175749 | 5.91 |
| GO_biological_processes | GO:0045766~positive regulation of angiogenesis | 0.029031787 | 2.31 |
| GO_biological_processes | GO:0050727~regulation of inflammatory response | 0.030367785 | 2.96 |
| KEGG_pathway | hsa04975:Fat digestion and absorption | 0.030429688 | 3.37 |
| KEGG_pathway | hsa00650:Butanoate metabolism | 0.032142023 | 4.06 |
| GO_biological_processes | GO:0006814~sodium ion transport | 0.032219408 | 2.63 |
| GO_biological_processes | GO:0048260~positive regulation of receptor-mediated endocytosis | 0.032549122 | 5.60 |
| GO_biological_processes | GO:0030198~extracellular matrix organization | 0.033745694 | 1.90 |
| GO_biological_processes | GO:0090181~regulation of cholesterol metabolic process | 0.033860949 | 9.98 |
| KEGG_pathway | hsa00460:Cyanoamino acid metabolism | 0.037147063 | 9.40 |
| GO_biological_processes | GO:0019370~leukotriene biosynthetic process | 0.037254292 | 5.32 |
| GO_biological_processes | GO:0006096~glycolytic process | 0.037352177 | 3.91 |
| GO_biological_processes | GO:0007626~locomotory behavior | 0.038188298 | 2.53 |
| GO_biological_processes | GO:0006935~chemotaxis | 0.040165559 | 2.18 |
| KEGG_pathway | hsa00591:Linoleic acid metabolism | 0.040543115 | 3.78 |
| GO_biological_processes | GO:0006641~triglyceride metabolic process | 0.040965189 | 3.80 |
| GO_biological_processes | GO:0006865~amino acid transport | 0.040965189 | 3.80 |
| GO_biological_processes | GO:0008203~cholesterol metabolic process | 0.041920546 | 2.74 |
| GO_biological_processes | GO:0009749~response to glucose | 0.041920546 | 2.74 |
| GO_biological_processes | GO:0019835~cytolysis | 0.042286818 | 5.07 |
| GO_biological_processes | GO:0016125~sterol metabolic process | 0.042286818 | 5.07 |
| GO_biological_processes | GO:0006544~glycine metabolic process | 0.042470726 | 8.87 |
| GO_biological_processes | GO:0007614~short-term memory | 0.042470726 | 8.87 |
| GO_biological_processes | GO:0072488~ammonium transmembrane transport | 0.042470726 | 8.87 |
| GO_biological_processes | GO:0090331~negative regulation of platelet aggregation | 0.042470726 | 8.87 |
| GO_biological_processes | GO:0070327~thyroid hormone transport | 0.042470726 | 8.87 |
| GO_biological_processes | GO:0045471~response to ethanol | 0.043775439 | 2.28 |
| GO_biological_processes | GO:0035690~cellular response to drug | 0.044520986 | 2.70 |
| GO_biological_processes | GO:0071333~cellular response to glucose stimulus | 0.044713088 | 3.07 |
| GO_biological_processes | GO:0009409~response to cold | 0.044769293 | 3.70 |
| GO_biological_processes | GO:0006956~complement activation | 0.044839931 | 2.45 |
| KEGG_pathway | hsa00140:Steroid hormone biosynthesis | 0.046805244 | 2.65 |
| KEGG_pathway | hsa00561:Glycerolipid metabolism | 0.046805244 | 2.65 |
| GO_biological_processes | GO:0006954~inflammatory response | 0.049012626 | 1.54 |

